# Phenology and flowering overlap drive specialization in pollinator networks

**DOI:** 10.1101/2020.09.08.287946

**Authors:** Paul Glaum, Thomas J. Wood, Jonathan R. Morris, Fernanda S. Valdovinos

## Abstract

Variation in diet breadth and specialization stems from fundamental interactions species have with their environment^1-3^. Consequently, understanding the drivers of this variation is key to understanding ecological and evolutionary processes, and will facilitate the development of predictive tools as ecological networks respond to environmental change^4,5^. Diet breadth in wild bees has been an area of focus due to both their close mutualistic dependence on plants, and because both groups are under threat from global biodiversity loss^6^. Though many of the principles governing specialization for pollinators have been identified^7,8^, they remain largely unvalidated. Using mechanistic models of adaptive foraging in pollinators^9,10^, we show that while temporal resource overlap has little impact on specialization in pollinators with extended flight periods, reduced overlap increases specialization as pollinator flight periods decrease. These results are corroborated empirically using pollen load data taken from bees with shorter (genus *Andrena*) and longer (genus *Lasioglossum*) flight periods across environments with both high and low temporal resource overlap. This approach reveals how interacting phenologies structure plant-pollinator networks and drive pollinator diet breadth via the temporal overlap of floral resources.

---

The dietary specialization of an organism sits at the nexus of numerous fundamental processes in ecology and evolutionary biology. Diet breadth helps define organismal niches^**1**^, mediates migration and dispersal potential^**2,11**^, shapes inter- and intraspecific competition^**3**,**12**^, influences species persistence in the face of environmental disturbance^**4**^, and can affect rates of speciation and extinction^**5**^. Even the simple assembly of the collection of dietary ranges of organisms in a community defines one of the major concepts in ecology, the food web. Therefore, identifying the drivers influencing diet breadth and specialization provides fundamental steps towards understanding a multitude of essential biological questions.

Much of the progress in understanding specialization in consumers comes from studies of herbivory^**1**^, particularly on insects^**13**^. Using wild bees and plant-pollinator networks as our study system, we extend the scope of those studies to include mutualistic interactions. A mutualism is a ubiquitous ecological interaction in which participant species benefit each other^**14**^, and pollination therefore differs from other insect herbivory because of the reproductive benefit offered to both consumer and consumed species. Bees depend on removing pollen from flowers to rear their offspring, providing a pollination service at the same time^**15**^. Pollination is also of particular interest given its role in supporting terrestrial biodiversity^**6**^ and agricultural output^**16**^, a role threatened by widespread declines in both pollinators and insect pollinated plants^**6**,**17**^.

Research on insect herbivores has largely focused on how adaptation to plant traits and defenses drives consumer concentration on single plant families^**18**,**19**^. However, other drivers of diet breadth have been identified (ref **7,8**, see Table S1). Here, we expand on the fundamental effect of resource density on a consumer’s diet breadth and specialization^**7**^, not in a spatial context, but temporally. This resource density is important in its interaction with consumer phenology. For long-lived organisms (e.g., predatory vertebrates), opportunities to access a single resource type principally depend on the resource’s spatial distribution. If a single resource type is physically dense, then a predator can specialize on it without needing to turn to alternatives, whereas if it is patchily distributed then a strategy of specialization is less optimal. However, for a comparatively short-lived organism like a bee whose adult flight period can be measured in days, the distribution of a resource type in time is the principal determinant of accessibility. A plant species that blooms for 30 days is *de facto* dense, continuously available resource for a bee species whose flight period lasts for only 25 days, but sparse and patchily distributed for a species which flies for 90 days. The temporal pattern of flowering can therefore be considered the functional equivalent of resource density for short-lived organisms. These ideas have proven influential but require concrete theoretical and empirical validation to solidify their conclusions.

Expanding upon this idea of temporal density, we hypothesize that the degree of temporal overlap in the availability of different resources (flowering overlap among co-occurring plant species) in combination with the fundamental life history phenologies of bees (adult flight period) determines realized diet breadth. Applied to plant-pollinator systems, we hypothesize that reduced flowering overlap will reduce the diet breadth of short-lived but not of long-lived pollinators (Fig. 1). Shorter-lived pollinators will experience markedly different resource availability based on the degree of temporal overlap of co-flowering species. High flowering overlap will allow shorter-lived pollinators a greater number of options while low overlap will restrict options. Longer-lived pollinators will be less constricted by the temporal overlap of co-occurring resources, often having equal number of options regardless of the degree of overlap (Fig. 1). Importantly, our hypothesis does not indicate that flight period will necessarily correspond with being more or less specialized. Instead, it describes how temporal resource overlap will affect diet breadth as a function of a pollinator’s flight period.

**Figure 1:**
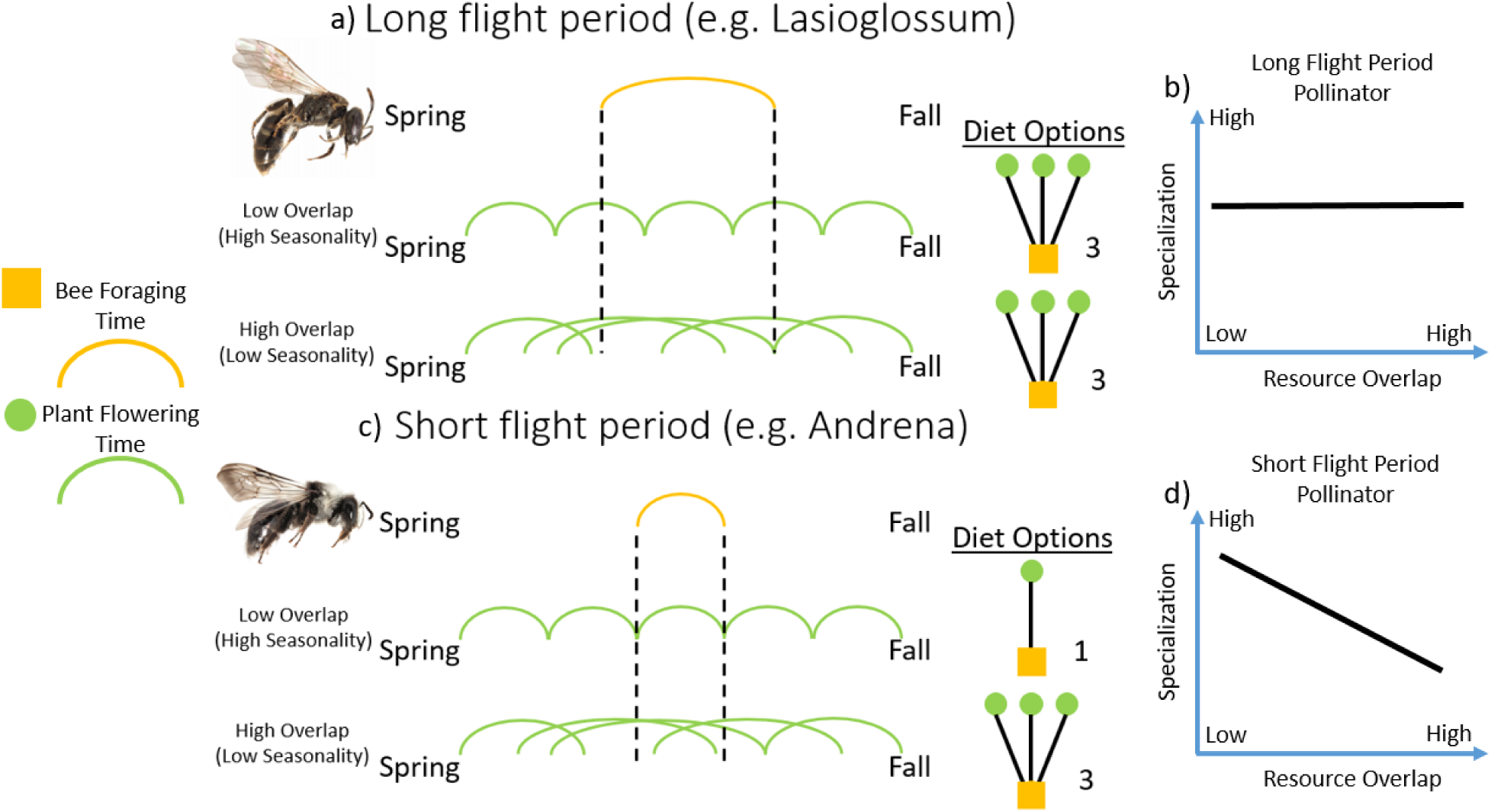
Graphical diagram of our hypothesis. Our hypothesis stems from the temporal overlap of flowering resources available to pollinators. Highly seasonal delimitations between different flowering populations create low levels of temporal overlap amongst different flowering resources. On the other hand, fewer seasonal delimitations between different flowering populations create higher levels of temporal overlap amongst different flowering resources. a) In long flight-period pollinators (e.g. *Lasioglossum*), differing levels of temporal resource overlap have limited to no effect on the degree of diet options available to the pollinator. b) This is predicted to lead to a non-significant relationship between overlap and specialization in long flight period pollinators. c) In short flight-period pollinators (e.g. *Andrena*), low and high temporal resource overlap can limit or expand pollinator diet options respectively. d) This is predicted to lead to a significant relationship between overlap and specialization in short flight-period pollinators.

We employed a multi-faceted approach to test our hypothesis. First, we leverage advances in modeling adaptive foraging in ecological networks^**9**,**10**^ to directly address the phenological mechanism driving diet breadth across different pollinator flight periods. Second, we utilize our intercontinental pollen load dataset of shorter lived *Andrena* bees and longer lived *Lasioglossum* bees from both a highly seasonal and a less seasonal environment which function as low and high temporal resource overlap treatments, respectively, to empirically validate our model outputs.

## Results

### Overview

Plant-pollinator dynamics are modeled using a dynamic consumer-resource approach that incorporates adaptive foraging of pollinators to mechanistically model pollinators’ consumption of floral rewards and reproductive services to plant species^**9**,**10**^ (see Supplementary Methods Table S2). The model’s adaptive foraging mechanism serves as a useful tool to implement and measure changing pollinator diet breadth (see Supplementary Methods). Phenology is integrated into the base model through modified sinusoidal wave functions which produce unique phenologies for every plant 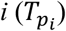 and animal pollinator 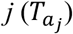. These functions 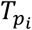 and 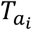 regulate floral reward production, adaptive foraging rates, pollinator visitation rates, and consequently, plant and pollinator reproduction across time (Tables S3-S4). The malleability of our phenology functions facilitates direct control over the availability of specific resources and the activity patterns of the animal pollinators over time in simulations (see Supplementary Methods, Fig S1-S5). The quantitative degree of overlap in co-occurring floral resources can be measured as the Total Resource Overlap (TRO) of the entire plant community, and the Averaged Resource Overlap (ARO) per plant species. Activity patterns of animal pollinators are set up to produce a range of different flight period lengths separated by differing lengths in between flight periods (Fig S6). Simulations use three fully connected bipartite network frameworks across 2072 combinations of plant and pollinator phenologies to produce 62160 unique networks used to model over 22 million plant-pollinator interactions (see Supplementary Methods).

Our pollen load data (see Supplementary Methods) comprises a large sample set of two bee genera, *Andrena* (Andrenidae) and *Lasioglossum* (Halictidae), collected both in the state of Michigan, USA (Nearctic) and the United Kingdom (Palearctic). Restricting our geographic scale to these two specific Holarctic locations aids in our ability to compare across both bee and plant communities due to their shared evolutionary history and overall similarity (see Supplementary Methods). The highly seasonal continental climate of Michigan^**20**^ produces lower degrees of overlap and highly seasonal flowering communities, while the more mild oceanic climate of the British Isles^**20**^ and its consistent temperatures produces longer, overlapping flowering times among plant species^**21**^.

Diet breadth or the degree of specialization in pollinators is measured both in-model with output on pollinator foraging effort and empirically with relative abundance of different pollen in bee pollen loads. We used two metrics that were applicable to both theoretical and empirical results for better direct comparisons, because simply counting unique taxonomic groups in the pollen data will not be applicable to theoretical results. We developed the first metric and labeled it Deviation from Generalism (DFG) (see Supplementary Methods, Table S5). Briefly, DFG is the normalized summation of all pairwise differences of pollinator foraging effort (or relative pollen load) on potential floral resources (see Supplementary Methods, Table S5). It ranges from 0 (perfect generalist) to 1 (perfect specialist). The second metric was the Coefficient of Variation (CV) of a pollinator’s foraging effort (or pollen load). Higher CV values occur with outlying values indicating specialization. With model results, both DFG and CV were applied to final foraging levels at the end of simulations as well as the average foraging levels across the last 1000 time steps (see Supplementary Methods, Fig S7).

### Model Results

Our simulation results (Figs. 2, 3) support our hypothesis (Fig. 1) demonstrating strong effects of temporal resource overlap on the diet breadth of short-but not long-lived pollinators. Fig. 2 shows the results of one network as an illustration of how resource overlap explains large portions of the variation in diet breadth of pollinators with shorter flight periods (Fig. 2a, c), but little for the diet breadth of pollinators with longer flight periods (Fig. 2b, d). The transition from high explanatory power to low explanatory power is apparent as pollinator flight period increases regardless if specialization was measured at the end of simulations or averaged across the last 1000 model time steps (Fig 3). This result is also consistent across varying proportional lengths of time between pollinator flight periods using either the DFG or CV metric across all networks tested (Fig S8-Fig S10). As predicted, low levels of resource overlap drive pollinators with short flight periods to specialize because potential resource options are limited at any given moment in time. As temporal resource overlap increases, potential options increase resulting in a more generalized diet breadth (Fig. 2a, c). Long-lived bees, on the other hand, do not experience the same limitation of potential diet options due to temporal resource overlap, because they are active most of the entire flowering season and can potentially access most or all of the flowering plants. Results were consistent regardless of whether resource overlap was measured as total resource overlap (TRO) or average resource overlap (ARO).

**Figure 2:**
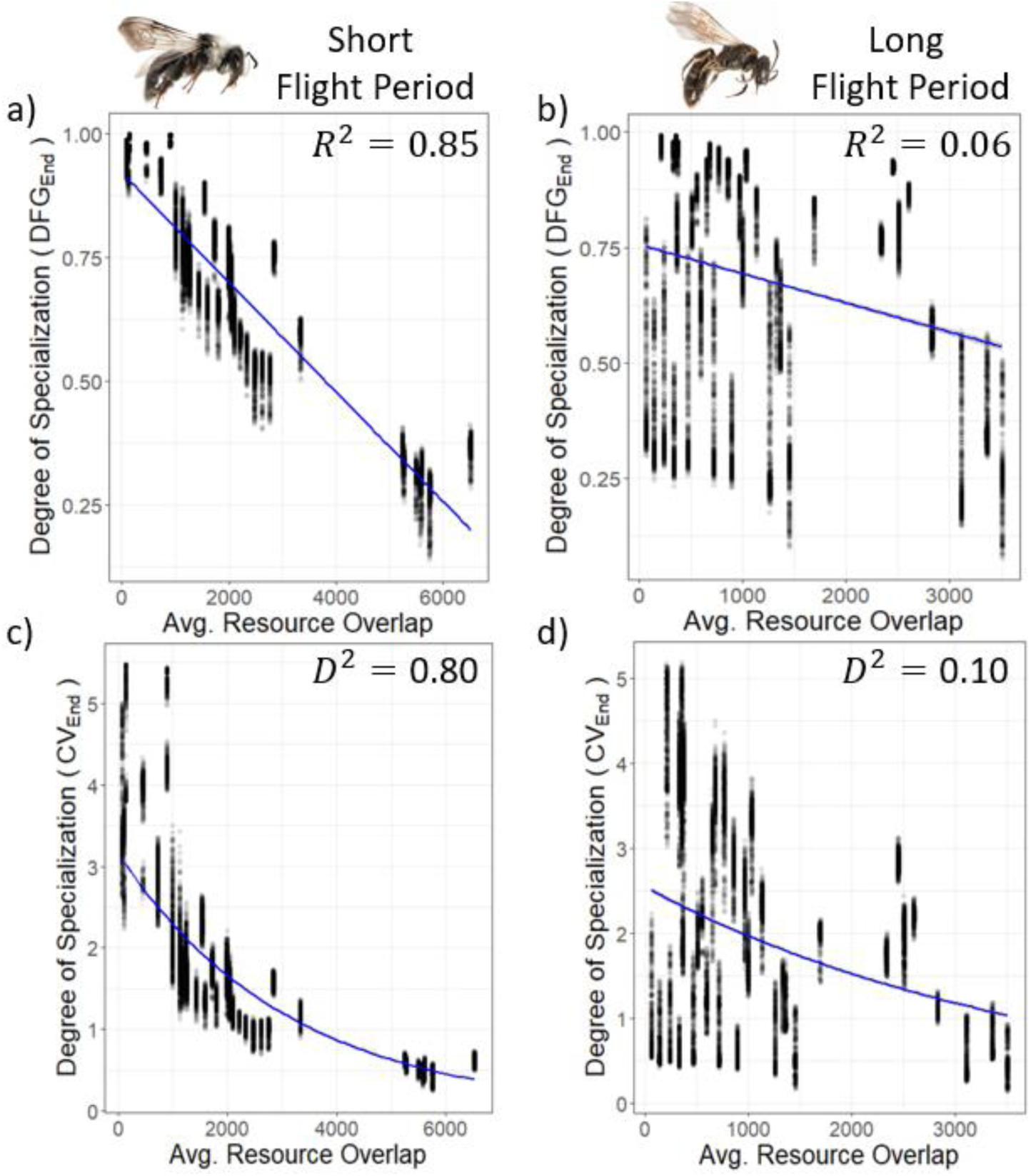
Temporal resource overlap drives specialization more strongly in short-lived than in long-lived pollinators. Model output on pollinator specialization taken across average plant overlap (ARO) for short flight period (flight period = 20 time steps) and long flight period pollinators (flight period = 300 time steps). Each dot represents the diet breadth metric of a single pollinator population based on their foraging effort per plant species. Diet breadth was measured at the end of simulations using **a)** DFG in a short flight period bee, F-statistic: 1.05e+05 on 1 and 18498 DF, p-value: < 2.2e-16, **b)** DFG in a long flight period bee, F-statistic: 1089 on 1 and 18498 DF, p-value: < 2.2e-16, **c)** CV in a short flight period bee, Residual deviance: 671.94 on 18498 degrees of freedom, **d)** CV in a long flight period bee, Residual deviance: 8244.4 on 18498 degrees of freedom. Example data taken from a network framework with 30 plant and 50 pollinator species. Higher values of both DFG and CV metrics indicate higher levels of specialization (narrower diet breadth) while lower values indicate a more generalized feeding range (wider diet breadth).

**Figure 3:**
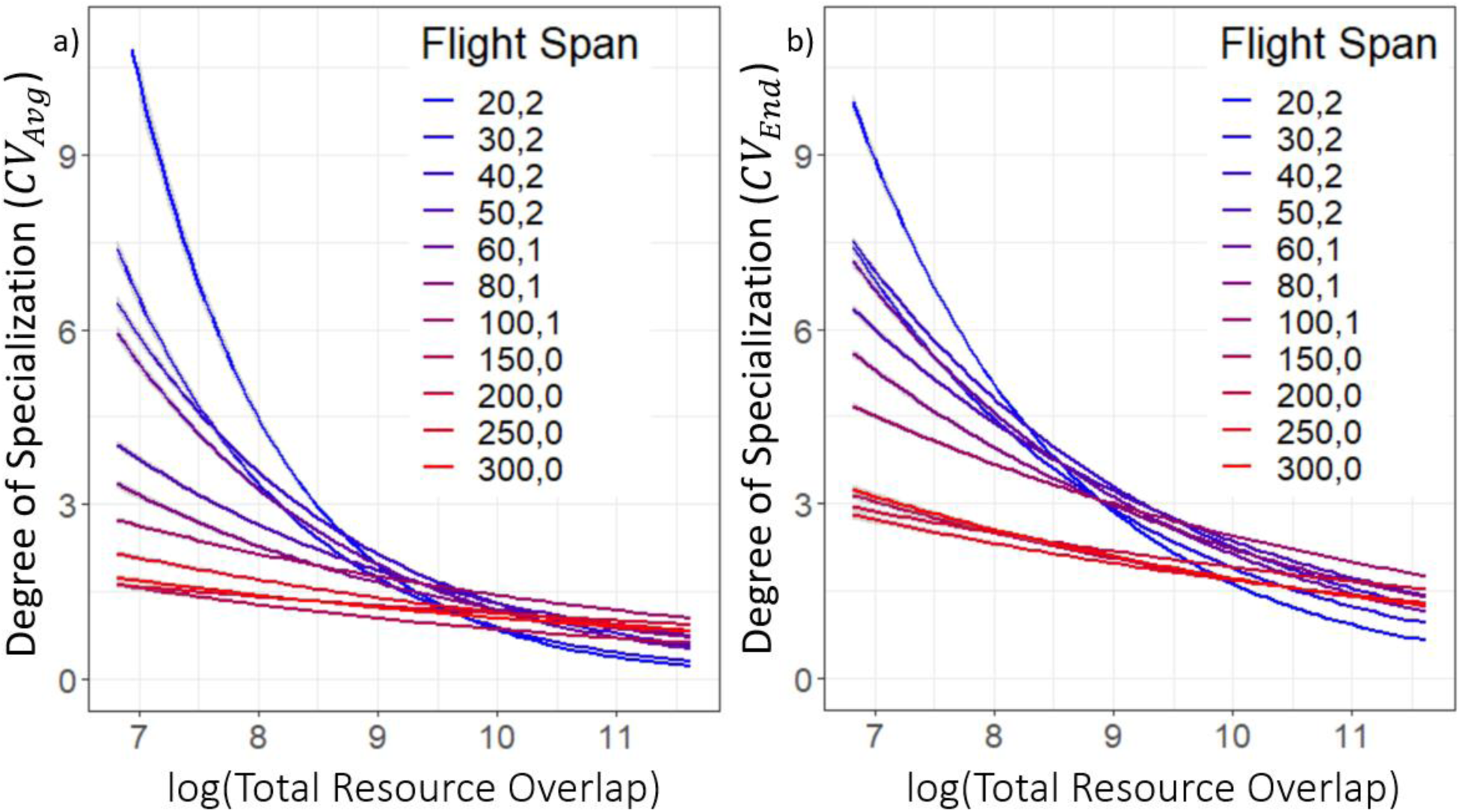
Interactive effect of temporal resource overlap and pollinator flight period on pollinator diet breadth. Representative sample of generalized linear models (Gamma distribution with log link functions) showing change in effect of Total Resource Overlap (TRO) on pollinator diet breadth in model simulations. Flight period labels indicate the length of the flight period and the number of phenology cycles between subsequent flight periods (see Supplementary Methods). Diet breadth shown as the degree of specialization measured by the CV of pollinator foraging effort per plant species taken **a)** at the end of simulations and **b)** averaged across the last 1000 time steps. Higher CV values correspond with a higher level of specialization (narrower diet breadth) while lower CV values indicate a more generalized feeding range (wider diet breadth). Example data taken from a network framework with 30 plant and 50 pollinator species.

### Pollen data results

Our empirical results support our hypothesis, showing more specialized diets of short-lived bees in more seasonal environments (Michigan) with lower flowering overlap, than in less seasonal environments (UK) with higher flowering overlap. Our pollen data from Michigan presents more single family specialists than the less seasonal UK dataset (Fig S11). Dietary specialization between bee genera and regions varied significantly at both the level of botanical family (χ^2^=27.1, p<0.001, Fig 4a) and genus (χ^2^=25.3, p<0.001, Fig 4b). In accordance with our hypothesis, there was an effect of region at the botanical family level, but only for the short-lived *Andrena* where species in Michigan were significantly more specialized than their British counterparts. In contrast, there were no differences for long-lived *Lasioglossum*. At the botanical genus level, the same trend was more strongly expressed, with *Andrena* in the UK showing a more generalized diet than their Michigan counterparts, but with no differences from *Lasioglossum* in either Michigan or the UK. Results were consistent when analyzed using the DFG metric (Fig S12).

**Figure 4:**
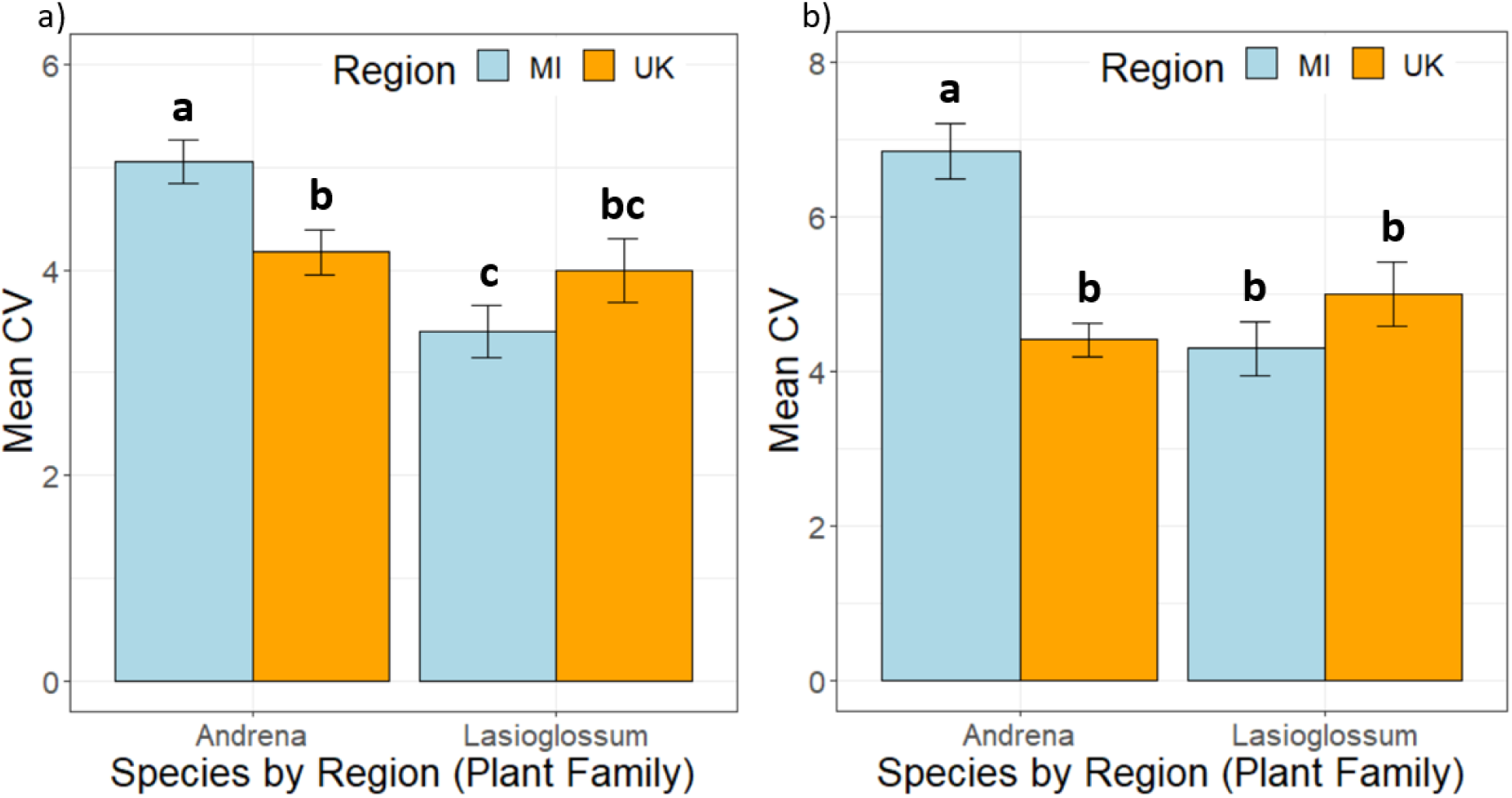
Empirical measures of dietary specialization. Short-lived *Andrena* bees show higher levels of specialization in highly seasonal (Michigan) compared to the weakly seasonal (UK), with no differences for long-lived *Lasioglossum* bees using the CV metric. Moreover, only in Michigan is there a difference in specialization between the two bee genera, showing a minimal impact of flight period in weakly seasonal environments. Effects are consistent at both the (a) botanical family and (b) botanical genus level. Bars headed by different letters are significantly different (p<0.05). Error bars show standard error.

## Discussion

Our model results mechanistically demonstrate how differing phenologies interact with varying degrees of temporal resource overlap to alter pollinator diet breadth. These results are corroborated through our regional pollen load data which also presents the first empirical validation of benchmark theories on adaptive foraging by consumers in general^**7**^ and pollinators in particular^**8**^. Furthermore, our results suggest broader implications regarding the drivers of bee diet breadth and diversification.

Global patterns of herbivory, particularly in insects, show strong trends of specialization at the botanical family level^**13**^. These patterns have been linked to phytochemical diversity^**19,22,23**^ which consequently drives an increase of both dietary specialization and species diversification in many herbivorous insect groups across a latitudinal gradient towards the tropics, most clearly seen and well-studied in Lepidoptera^**22**,**24**^. However, bees break sharply from this pattern, with a relatively low diversity in the tropics when compared to Mediterranean and xeric environments^**8,25,26**^. Tropical environments are dominated by highly generalized, often social species that visit a huge variety of botanical families^**27**,**28**^. Tropical areas have the highest levels of global phytochemical diversity, but their flowering patterns have wider phenological variation than temperate areas at both the inter- and intraspecific levels. This results in tropical floral communities that *de facto* show higher resource overlap in any particular year^**29**,**30**^ than temperate communities. Our theoretical and empirical results suggest that bees’ strong departure from the typical global pattern in herbivory^**13**^ may be due to a relatively stronger effect of phenology on their diet breadth than the phytochemical restrictions suggested for other insect herbivores. In fact, tropical bees have not evolved anything approaching the same high degree of dietary specialization as seen in the tropical Lepidoptera despite existing in the same extremely phytochemically diverse landscapes. For further discussion of this point, see Supplementary Discussion.

Our modeling developments present a flexible operational basis going forward. Integration of time dependent functional components into species behavior and traits show that even high dimensional network models can be further expanded in tractable ways to include the dynamics of organismal phenology. Additionally, our metrics for diet breadth complement each other (see Supplementary Methods) and reflect past methods (Fig S10). These metrics also integrated well with empirical pollen load data sets that can be used to vet theoretical predictions. By expanding pollen datasets globally and incorporating empirically vetted plant-pollinator phenology, future research can address how foraging competition occurs across overlapping phenologies and what effects changing climatic conditions can have on network stability.

## Supporting information

Supplementary Methods and Supplementary Results

Table S9 - Pollen load data

Table S10 - Phenology parameters

## Acknowledgements

Part of this project was supported by the National Science Foundation grant DEB-1834497 awarded to F.S.V.

## Author Contributions

F.S.V. conceived the project. T.J.W., P.G., F.S.V., and J.R.M. developed the conceptual basis for the project. T.J.W. identified suitable collection sites and bee genera for the empirical study design, collected bee and pollen data, and identified specimens. P.G. and F.S.V. developed dynamic model, simulation design, code, and diet breadth metrics. P.G. implemented simulations and analyses. T.J.W. and P.G. wrote the first draft, and all authors edited and revised the manuscript.

## Supplementary Information

Supplementary Methods, Supplementary Discussion, Supplementary Figures and Tables.

## Data Availability

Pollen load data is available as Supplementary Table S9.

## Code Availability

Simulation code and simulation data are available at the repository: https://github.com/fsvaldovinos/Phenology. Phenology parameters used in the simulation portion of our study are available in Table S10.

