## Supplementary Methods and Supplementary Results for "Phenology and flowering overlap drive specialization in pollinator networks"

*Historical Background*

Beyond the important role of the phytochemical landscape driving insect herbivore diet breadth, herbivorous insects are subject to a list of general factors that affect the diet breadth of foragers. Originally broadly conceptualized for hunting animals by MacArthur & Pianka[7], this list was adapted for application to a pollinator diet by Wasser et al[8]. These factors are listed in Table S1 with common factors between the two treatments.

**Table S1:** Factors favoring specialization in foraging as detailed in MacArthur & Pianka[7] and their corresponding construction in Waser et al's[8] study on pollinator specialization. The third column describes the common factor in each treatment. The factor noted in green highlights the work which helped inspire our study.

| Factors favouring specialisation of diet for hunting animals (MacArthur and Pianka 1966) | Factors favouring generalisation of diet for pollinators (Waser et al. 1996) | Common factor |
|---|---|---|
| **Greater food density** | **Pollinator lifespan is long relative to the flowering of individual plant species** | **The density of resources, as measured relative to the foraging range/period of the animal** |
| Increased mobility of animal, or decreased environmental resistance to movement | Travel is costly | The cost associated with travelling to many or few resources |
| Increased differences between prey types, or increased specialisation of pursuing behaviour | Floral rewards are similar across plant species | The inherent differences in resource quality |
| Increased mobility of prey, or greater difficulty in pursuit | Constraints of behaviour and morphology are minor | The inherent difficulty involved in accessing different resources |

*Baseline model*

As with recent studies of ecological networks[10 31], the makeup of the overall network model used here has two major fundamental components: the structure of the pollination networks themselves and the dynamics occurring on the networks.

The structure of a network describes which links (representing species interactions) between plants ($p$) and animal pollinators ($a$) are present or absent, regardless of the strength of the link. Typical ecological network studies have a collection of unique topological connections across different networks, meaning only certain links of the possible links between plants and pollinators can potentially be realized

(Fig S1a). However, the networks we used in the models/simulations here were fully connected, meaning all connections are possible (Fig S1b). The baseline of these networks in our study are sourced from an empirical pollination network[32]. Using a real-world pollination network as a basis ensured a plausible ratio of flowering plant populations to pollinator populations, which in this case was 58 plant species to 100 pollinator species. The network was then fully connected (see Figure S1) such that each pollinator had potential access to all plant species and vis-versa. Fully connecting the plant-pollinator network was done in order to give each pollinator population the maximum within-model range of dietary options. Doing so allows us to test the ability of the phenological mechanism to drive plasticity in pollinator specialization without a priori constraints on pollinator behavior and network structure. The size of this basic network framework was then modified to test effects of species richness, resulting in 3 network size classes ($N_p$=plant richness, $N_a$=animal pollinator richness): $N_p = 58$ & $N_a = 100$, $N_p = 48$ & $N_a = 71$, and $N_p = 30$ plants & $N_a = 50$. While only these 3 network frameworks were used, the exact topology of each changed with thousands of permutations of phenological parameters (see below).

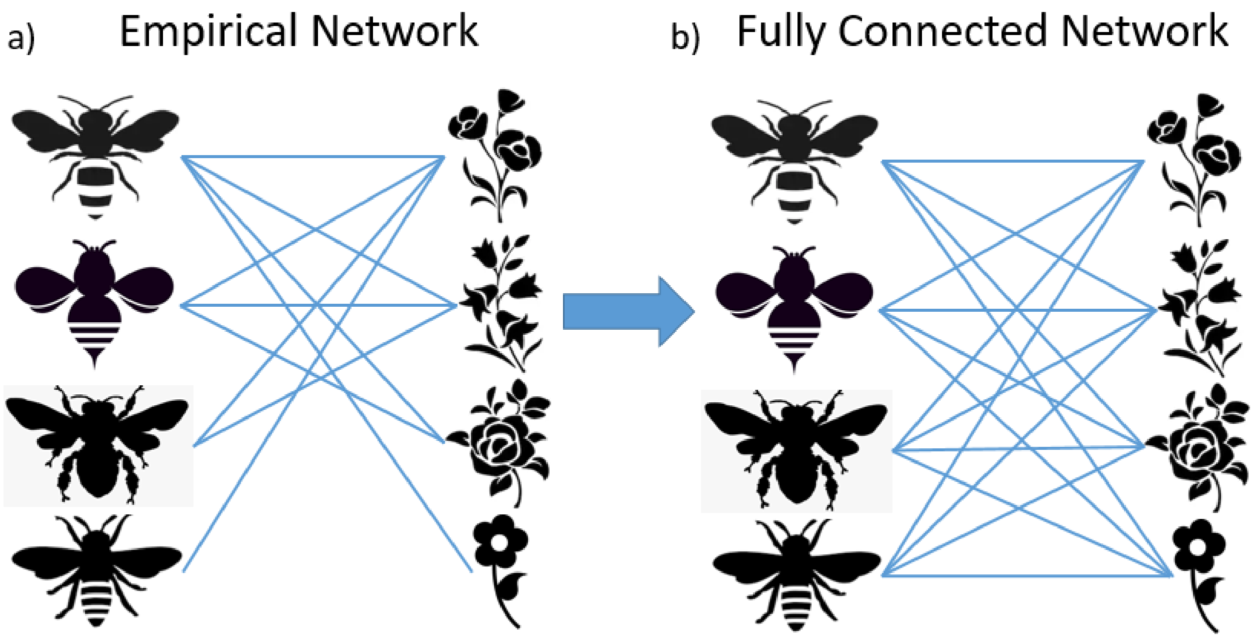


**Figure S1: Underlying network structure.** Example diagram describing the creation of the baseline network structure. Pictures of bees represent populations of pollinators and pictures of flowering plants represent populations of flowering plants. Blue links between them represent a potential pollination interaction between plants and pollinators. a) Example of a starting empirical network. b) Example of fully connecting the empirical network to use in simulations.

The dynamics occurring on the networks refer to the internal population demographics (birth and death) and species interactions taking place amongst the species in the network. In the case of our model, species interactions refer to competition for resources, consumption of resources, and pollination events. Dynamics were simulated based on the work of Valdovinos et al[9] which mechanistically modeled pollination as a consumer-resource interaction by separately accounting for vegetative abundance and floral rewards consumed by pollinators. The model tracks the adaptive dynamics of each plant species' population dynamics ($p_i$), each animal pollinator species population dynamics ($a_j$), each plant species' pool of floral rewards ($R_i$), and the adaptive dynamics of the per-capita foraging effort preferences of each pollinator species for each plant species ($\alpha_{ij}$). Model parameters are described in Table S2.

**Table S2:** Baseline model state variables and parameters. In Mean Value column, * indicate initial conditions, $k_{aj}$ is the number of interactions of animal *j*

| Definition | Symbol | Dimension | Mean Value |
|---|---|---|---|
| Density of plant population *i* | $p_i(t)$ | individuals area$^{-1}$ | 0.7* |
| Density of animal population *j* | $a_j(t)$ | individuals area$^{-1}$ | 0.7* |
| Density of floral resources of plant population *i* | $R_i(t)$ | mass area$^{-1}$ | 0.7* |
| Foraging Effort of pollinator *j* on plant *i* | $\alpha_{ij}(t)$ | none | $\frac{1}{k_{\alpha j}}$ * |
| Visitation Efficiency | $\tau_{ij}$ | visits area time$^{-1}$ individuals$^{-1}$ individuals$^{-1}$ | 1 |
| Expected number of seeds produced per pollination event | $e_{ij}$ | individuals visits$^{-1}$ | 0.8 |
| Per capita mortality rate of plants | $\mu_i^P$ | time$^{-1}$ | 0.002 |
| Conversion efficiency of floral resources to pollinator births | $c_{ij}$ | individuals mass$^{-1}$ | 0.2 |
| Per capita mortality rate for pollinators | $\mu_j^A$ | time$^{-1}$ | 0.003 |
| Pollinator extraction efficiency of resource $R_i$ in each visit | $b_{ij}$ | individuals visits$^{-1}$ | 0.4 |
| Max fraction of total seeds that recruit to plants | $g_i$ | none | 0.4 |
| Intra-specific competition coefficient for plants | $u_i$ | area individuals$^{-1}$ | 1.2 |
| Inter-specific competition coefficient for plants | $w_i$ | area individuals$^{-1}$ | 0.002 |
| Production rate of floral resources | $\beta_i$ | mass individuals$^{-1}$ time$^{-1}$ | 0.2 |
| Self-limitation parameter for resource production | $\phi_i$ | time$^{-1}$ | 0.04 |
| Adaptation rate of pollinator foraging effort | $G_j$ | none | 2 |

Changes in population density of plant species ($p_i$) are calculated through:

$$\frac{dp_i}{dt} = \gamma_i \sum_{j\in A_j} e_{ij}\sigma_{ij}V_{ij} - \mu_i^{p_i} p_i \qquad \text{Eq (1)}$$

The second term in the equation describes background mortality, where $\mu_i^p$ is the constant density-independent per-capita mortality rate of plant $i$.The first term describes plant population growth where $\gamma_i$ describes the realized fraction of seeds that successfully recruit to adults:

$$\gamma_i = g_i(1 - \sum_{l\neq i\in P} u_l p_l - w_i p_i) \qquad \text{Eq (2)}$$

where $g_i$ is the maximum fraction of seeds that can potentially recruit to fecund adulthood. The recruitment is subject to both interspecific ($u_l$) and intraspecific ($w_i$) competition with $u_l < w_i$. The parameter $e_i$ in Equation (1) is the constant max seed set induced by a pollination event between plant and pollinator. The quality of each pollination event is determined by the fraction of pollen from conspecific plants on a pollinator compared to other plant species pollen. This fraction is proportional to the visits each animal pollinator ($a_j$) makes on each plant species ($p_i$). We label this term, $\sigma_{ij}$ and define it as follows:

$$\sigma_{ij} = \frac{V_{ij}}{\sum_{k \in P_j} V_{kj}} \qquad \text{Eq (3)}$$

where $V_{ij}$ is the frequency of visits by animal species $j$ to plant species $i$ and it defined by:

$$V_{ij} = \alpha_{ij}\tau_{ij}T_{a_j}(t)a_j p_i \quad \text{Eq (4)}$$

Visits from pollinator $j$ to plant species $i$ are zero ($V_{ij} = 0$) if the two do not interact. Pollinator $j$'s visitation efficiency on plant $i$ is determined by the parameter $\tau_{ij}$ and is fixed at 1 for this study so as not to bias any pollination interaction over others and affect visitation preferences a priori. The function $T_{a_j}(t)$ is the phenological determinant of activity of pollinator $a_j$, in this case controlling the flight period of $a_j$. See Phenology section below for more. The dimensionless function $0 \leq \alpha_{ij} \leq 1$ is the foraging preference of pollinator $j$ on plant $i$ and changes over time as defined by:

$$\frac{d\alpha_{ij}}{dt} = T_{a_j}(t)G_j\alpha_{ij}\left(c_{ij}\tau_{ij}b_{ij}R_i - \sum_{k \in P_j}\alpha_{kj}c_{kj}\tau_{kj}b_{kj}R_k\right) \qquad \text{Eq (5)}$$

where $G_j$ is the basal adaptation rate of foraging preference. Higher or lower rates of $G_j$ produce faster or slower rates of adaptation based on changes seen inside the parentheses of Eq (5). $c_{ij}$ represents the constant per-capita conversion efficiency of pollinator $j$ converting plant $i$'s floral resources into $j$'s births. $b_{ij}$ is the constant efficiency of pollinator $j$ extracting plant $i$'s floral resources ($R_i$). Pollinator $j$ allocates more foraging effort to plant species $i$ whenever such reallocation increases $j$'s food intake. Such reallocation causes a commensurate reduce in foraging effort from other plant species. For every animal pollinator species $j$, $\sum \alpha_{ij} = 1$ for all plant species. Finally, the function $T_{a_j}(t)$ appears here to limit the adaptation of foraging preference only to periods when pollinators are actively flying.

Each plant $i$'s floral resources, $R_i$, changes over time as defined by:

$$\frac{dR_i}{dt} = T_{p_i}(t)\beta_i p_i - \phi_i R_i - \sum_{j \in A} V_{ij} b_{ij} \left(\frac{R_i}{p_i}\right) \qquad \text{Eq (6)}$$

where $\beta_i$ is plant $i$'s per-capita resource production rate and $\phi_i$ is a constant self-limitation parameter. Rewards of plant $i$ are removed with an efficiency $b_{ij}$ by pollinator $j$ in proportion to the amount of visits, $V_{ij}$. The function $T_{p_i}(t)$ controls the phenological expression of resource production in each flowering plant $i$. Further details are provided below in Phenology section. The population dynamics of the animal pollinators are then defined by:

$$\frac{da_j}{dt} = \sum_{i \in P} c_{ij} V_{ij} b_{ij} \left(\frac{R_i}{p_i}\right) - \mu_j^A a_j \qquad \text{Eq (7)}$$

where $c_{ij}$, $V_{ij}$, and $b_{ij}$ are as defined above. Pollinator population growth is driven by the sum of resources gathered from pollination visits while death occurs at a constant rate $\mu_j^A$.

*Incorporating phenology into baseline model*

Creating temporal plasticity in network connectivity is implemented by modifying the rates controlling interactions between plants and pollinators in the network. In other words, the expression of certain species' rates/behaviors are modified by time dependent functions which activate and deactivate said rates across time. In flowering plant species, this is implemented by modifying the production of floral rewards ($R_i$) that drive pollinator foraging preference ($\alpha_{ij}$) and reproductive benefit for both plants and pollinators. In Eq (6), the rate of floral reward production ($\beta_i$) is regulated across time steps by the phenology function $T_{p_i}(t)$, Eq (8). Parameter definitions are given in Table S3.

$$T_{p_i}(t) = \left| f(b_{bv}) * \sin\left(b_{pt}\pi t - b_{w_s}\right) * \frac{\left(1 + square\left(\frac{b_{pt}}{2b_{bt}}\pi t - \frac{b_{w_s}}{b_{w_a}}, \frac{25}{b_{bt}}\right)\right)}{2} \right| \qquad \text{Eq (8)}$$

**Table S3:** Phenology parameter definitions for $T_{p_i}(t)$

| Phenology Parameter | Description | Definition |
|---|---|---|
| Bloom Period ($b_p$) | No. of time steps with floral resource production ($\beta > 0$) | Defined by user |
| Bloom Period translated ($b_{pt}$) | Transforming $b_p$ into useable form for $T_{p_i}$ | $\frac{1}{b_p}$ |
| Bloom break value ($b_{bv}$) | No. of cycles between subsequent blooms | Defined by user |
| Bloom Break Translated ($b_{bt}$) | Transforming $b_{bv}$ into useable form for $T_{p_i}$ | $\frac{b_{bv}}{2} + 0.5$ |
| ($d$) | % of maximum separation between $T_{p_i}$ | 1 |
| Separation distance ($p_{s_d}$) | Distance on the timeline between consecutive $T_{p_i}$, here $N_p$ is the number of plant species | $\frac{\left(\frac{2\pi b_{bv}}{N_p}\right)}{d}$ |
| Wave Shift ($b_{w_s}$) | Separation distance assigned to each plant species in sequence. | $\{p_{s_d} * n\}_{n=0}^{N_p-1}$ |
| Wave Augment ($b_{w_a}$) | Augment to save shift in the square wave | $b_{bv} + 1$ |
| $f(b_{bv})$ | Function mitigating lost resources with higher break value by increasing floral reward production | $b_{bv} + 2$ |

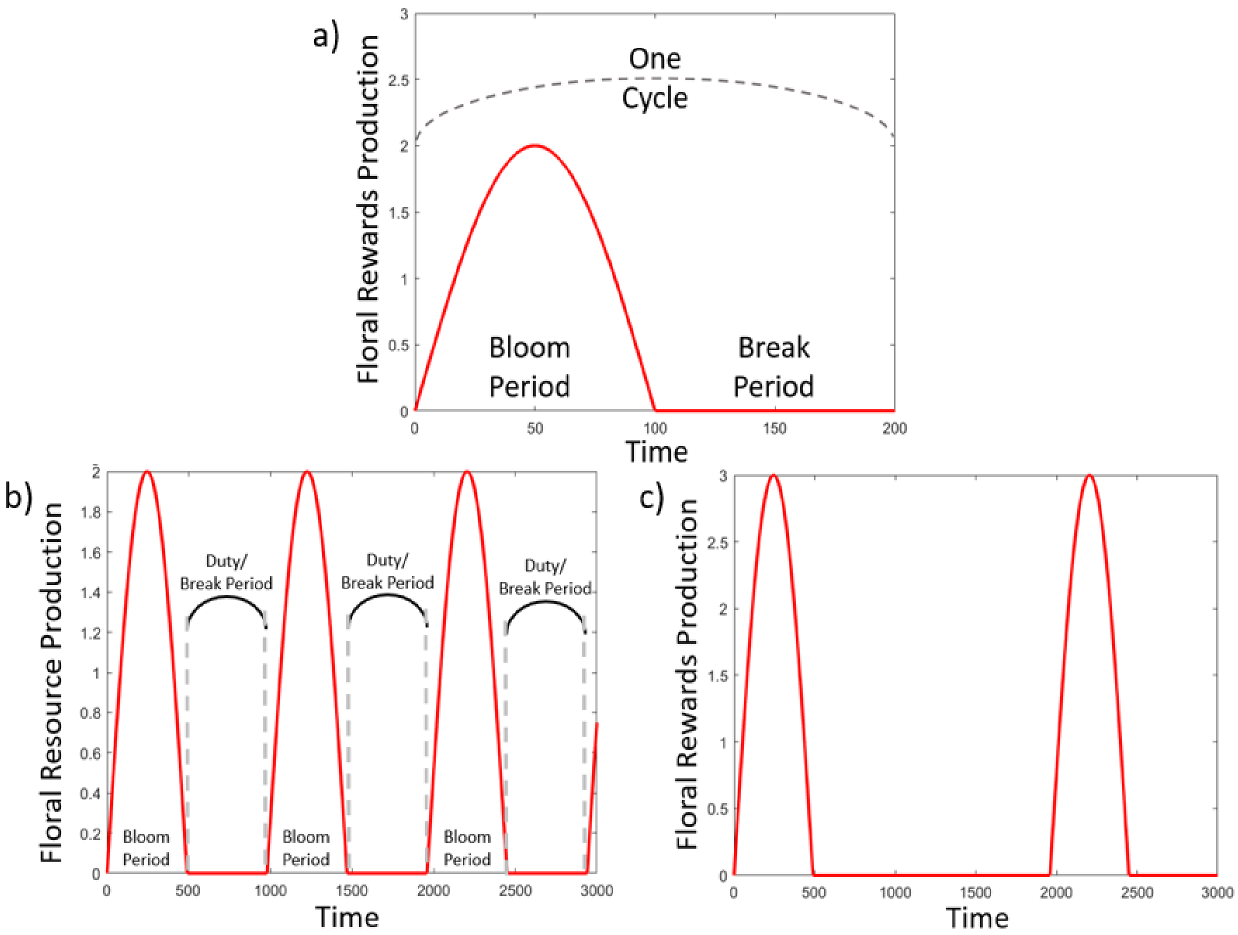


**Figure S2: Example $T_{p_i}$ figures.** A) Detail of the components of each $T_{p_i}$. B) Example of $T_{p_i}$ run across time. Bloom Period = 490, Break Period = 0. C) Example of $T_{p_i}$ run across time. Bloom Period = 490, Break Period = 1.

The form of $T_{p_i}(t)$ is a composite of sine and square waves which produce a continuous bounded function (always ≥ 0), with both active and inactive periods in one “cycle,” the bloom period and break period respectively. Bloom and break periods are controlled through the two parameters “bloom period” and “bloom break value” respectively (Fig S2a). The bloom period corresponds to the number of time steps that a plant will produce floral rewards from start to finish. Floral resource production follows a sinusoidal growth and decay during the bloom period, with peak production occurring during the middle of the bloom period (Fig S2a). Each bloom period is accompanied by a corresponding break period or “off period” where the value of $T_{p_i} = 0$. This is similar to the “duty” of a traditional square wave function. This break period is of equal length to the bloom period such that there is an even split between active and inactive parts of a single cycle when the break value is 0. In this default formulation, $T_{p_i}$ will cycle through active and inactive periods (Fig S2b), thereby modifying the production of floral rewards. When break value > 0, then $T_{p_i}$ will skip the number of cycles assigned to the break value before the ensuing bloom period (Fig S2c). For the purpose of parity in comparing across different bloom periods, the length of a cycle's break period scales with its bloom period. This facilitates normalizing available resource quantities between bloom period.

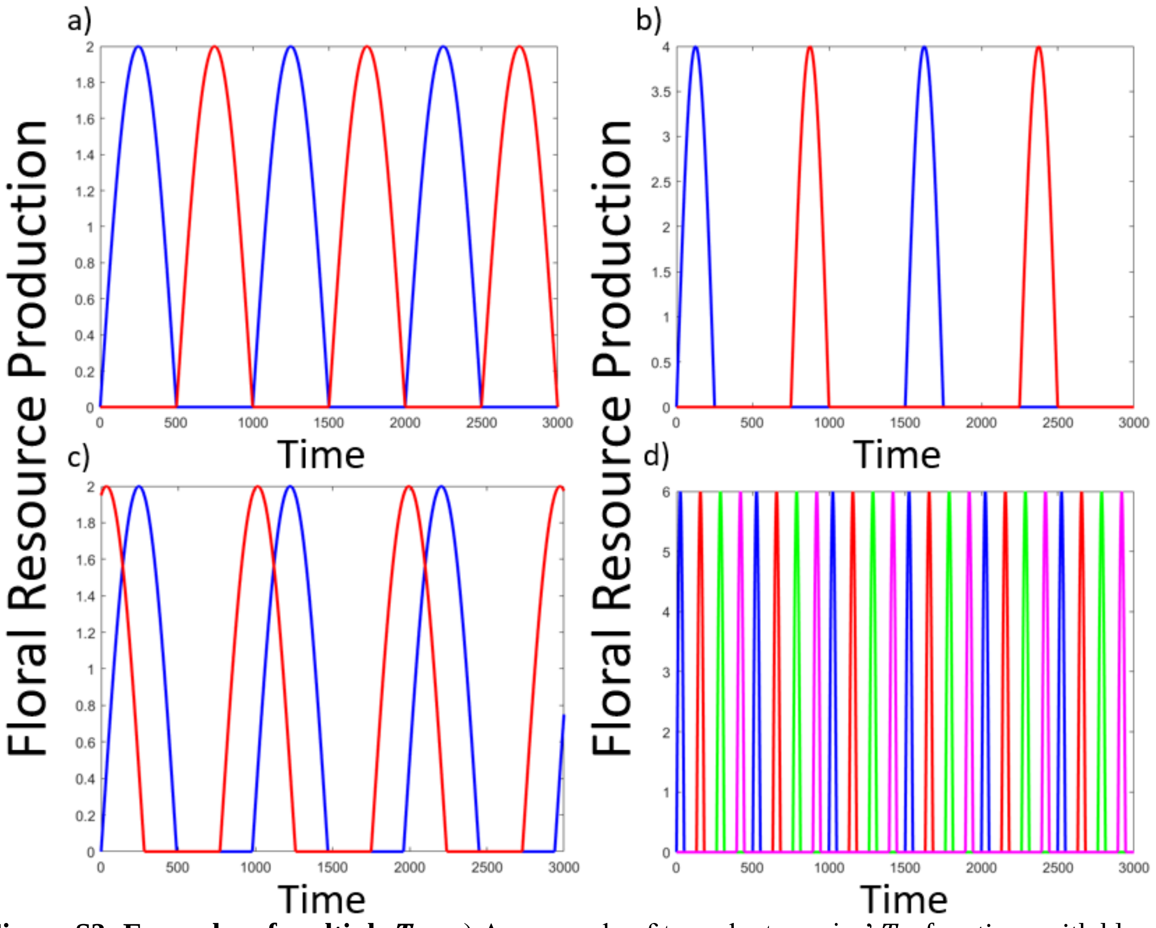


**Figure S3: Examples of multiple $T_{p_i}$.** a) An example of two plant species' $T_{p_i}$ functions with bloom period of 500 and break values of 0. b) An example of two plant species' $T_{p_i}$ functions with bloom periods of 250 and break values of 2. c) An example of two overlapping plant species' $T_{p_i}$ functions with bloom periods of 500 and break values of 0. d) Example of multiple $T_{p_i}$ for $i \in (1,4)$ with bloom periods of 125 and break value of 4.

Every $p_i$ is assigned its own $T_{p_i}(t)$ and their separation along the time of a simulation is put at a maximum (Fig S3a) in simulations, depending upon the break value (Fig S3b). Though the model does allow for more direct overlap (Fig S3c). This all then scales with the number of plant species, $p_i$, in the system (Fig S3d).

Phenology in bee species, $T_{a_i}$, is formulated similarly across visits ($V_{ij}$) in Eq (4) and $G_j$, the basal adaptation rate of foraging preference ($\alpha_{ij}$), in Eq (5). Both visitation and adaptation are modified by $T_{a_i}$ because foraging preference should not change when there is no active foraging. The formulation of $T_{a_i}$ is given in Eq (9) and its parameter definitions are given in Table S4. Similar to $T_{p_i}(t)$, overall control of the specific form of $T_{a_i}$ is governed by two parameters, "flight period" and "flight break value." The general form of $T_{p_i}(t)$ and $T_{a_i}(t)$ was chosen heuristically based on its ability to interface with the baseline model and the ability to maintain sufficient control over the specific form of the function with only two parameters. This makes it both readily applicable from a research stand point and relatively user friendly for other interested researchers.

$$T_{a_i}(t) = \left| f(f_{bv}) * \sin\left(f_{pt}\pi t - f_{w_s}\right) * \frac{\left(1+square\left(\frac{f_{pt}}{2f_{bt}}\pi t-\frac{f_{w_s}}{f_{w_a}},\frac{25}{f_{bt}}\right)\right)}{2} \right| \quad \text{Eq (9)}$$

**Table S4:** Phenology parameter definitions for $T_{a_i}(t)$

| Phenology Parameter | Description | Definition |
| --- | --- | --- |
| Flight Period ($f_p$) | No. of time steps with active flight ($V > 0$ & $G_j > 0$) | Defined by user |
| Flight Period translated ($f_{pt}$) | Transforming $f_p$ into useable form for $T_{a_i}$ | $\frac{1}{f_p}$ |
| Flight break value ($f_{bv}$) | No. of cycles between subsequent flights | Defined by user |
| Flight Break Translated ($b_{bt}$) | Transforming $f_{bv}$ into useable form for $T_{a_i}$ | $\frac{f_{bv}}{2} + 0.5$ |
| ($d$) | % of maximum separation between $T_{a_i}$ | 1 |
| Separation distance for animal phenology ($a_{s_d}$) | Distance on the timeline between consecutive $T_{a_i}$, here $N_a$ is the number of animal pollinator species | $\frac{\left(\frac{2\pi f_{bv}}{N_a}\right)}{d}$ |
| Wave Shift ($f_{w_s}$) | Separation distance assigned to each animal species in sequence. | $\{a_{s_d} * n\}_{n=0}^{N_a-1}$ |
| Wave Augment ($f_{w_a}$) | Augment to save shift in the square wave | $f_{bv} + 1$ |
| $f(f_{bv})$ | Function mitigating lost flight time with higher break value by increasing peak flight activity | $f_{bv} + 2$ |

With both plant and animal phenology integrated into the model, the temporal plasticity of pollination interactions and network topology can be realized. When a pollinator $j$'s flight/foraging time overlaps with a plant $i$'s flowering time, the pollination interaction is potentially active (depending on foraging preference $\alpha_{ij}$) and counted as a "link" in the network for the duration of the overlap (Fig S4). Otherwise, the link is inactive and interactions between the plant and pollinator do not occur.

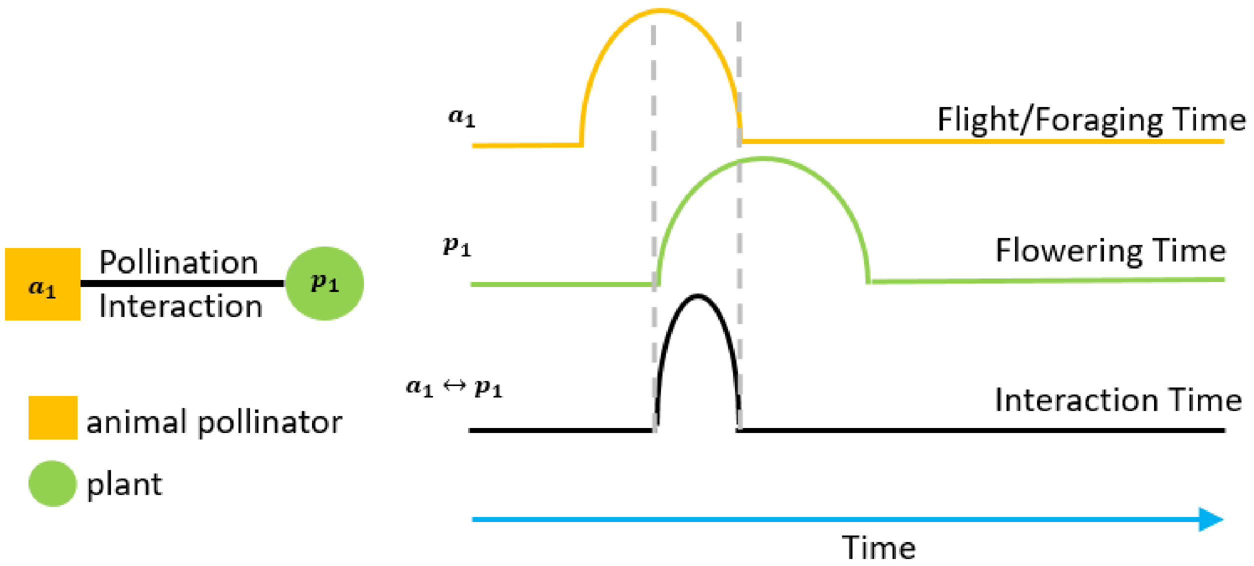


**Figure S4: Phenology controlled interaction.** Diagram of an example pollination interaction in time due to phenology of pollinator and plant. Note, while only one active period is shown for the pollinator and plant, this is only for readability in the diagram. In model simulations, all phenology functions cycle as shown in Fig S2&S3.

This framework scales up to multiple interactions and ultimately the network level as network topology changes dynamically as potential interactions activate and deactivate in the network across time. A diagram of this is provided in a small example plant-pollinator network is provided in Fig S5. Given foraging preference ($\alpha$), even a potentially active interaction may not be particularly active, but only during phenological overlap can the interaction potentially occur.

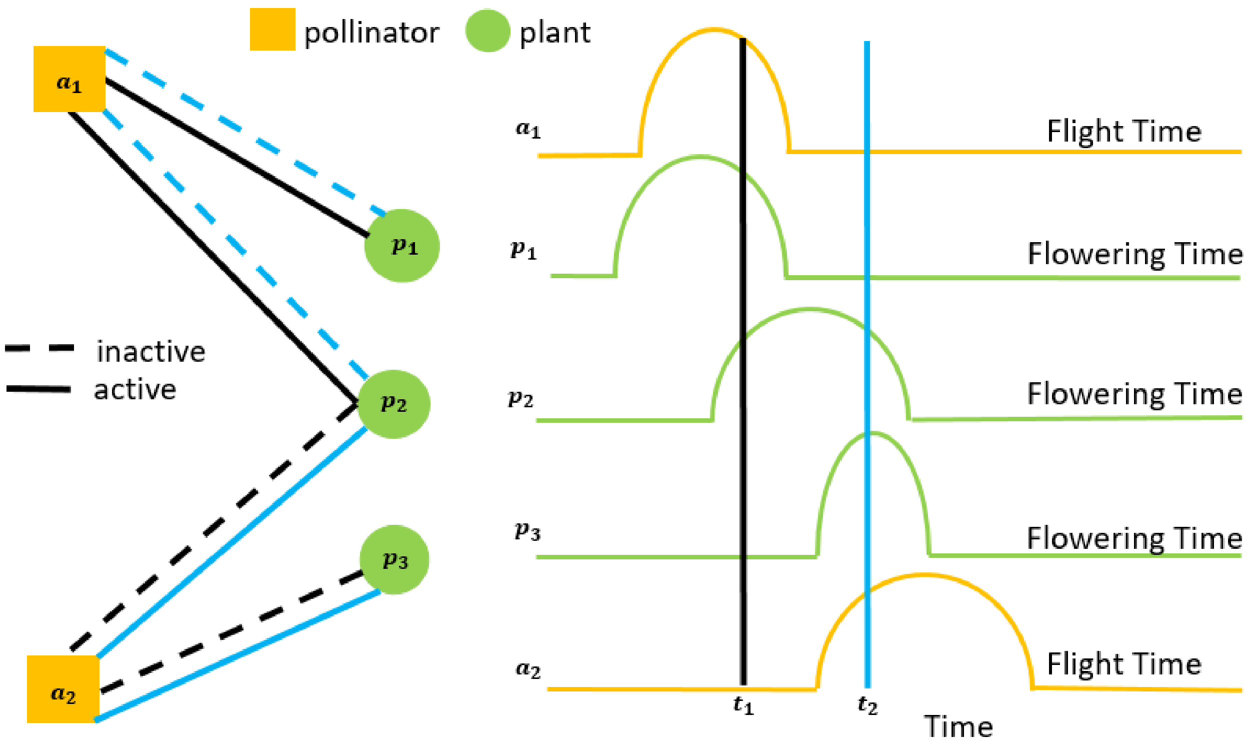


**Figure S5: Diagram of example network with phenology.** Example diagram of interacting phenologies changing the active topology of a small example network. At different times, as indicated by the black ($t_1$) and blue bars ($t_2$), different links in the network at active. This scales up to the networks used in simulations and analysis here. Note, while only one active period is shown for each pollinator and plant, this is only for readability in the diagram. In model simulations, all phenology functions cycle as shown in Fig S2&S3.

*Experimental simulation design*

Without phenology induced oscillations, baseline model simulations generally achieve reach asymptotic dynamic behavior in approximately 3000 time steps. In order to ensure that transient dynamics minimally affect measured model output, models here were run for 6000 time steps to reach asymptotic dynamics. No burn in period without phenology is used in simulations as that would produce preceding bias to foraging preference before phenology functions are activated. All plant and pollinator species experience their phenological changes in rates from the beginning of simulations.

Baseline parameters for plants (non-phenological), including competition coefficients, floral reward productivity, reward self-limitation, and mortality differ among plant species by assigning unique values stochastically varied within 0.1% of a mean value (Table S2). Competitive coefficients of plants are kept low ($e^{-3}$) in order to reduce effects on pollinator diet breadth through abundance loss and potential species loss in plants (Table S1). Our simulations intentionally produced no extinctions in plants or animals. Baseline parameters for animal pollinators (non-phenological, Table S2), including extraction efficiency, conversion efficiency, and mortality differ among pollinator species by assigning unique values stochastically varied within 0.5% of a mean value (Table S2). Similar to using fully connected networks, limiting variability in baseline parameter values across species and pollination interactions within single simulations allows us to focus analysis on the effect of phenology and temporal resource overlap on specialization. Distinct differences in productivity between plants or efficiencies in pollinators through more varied parameter values would bias foraging preferences towards certain interactions more than others. For example, a highly productive plant or a pollinator with uniquely high fundamental efficiency towards one plant species over others would drive specialization by assumption alone. Keeping parameter values closely centered around the mean for all possible interactions limits this bias while still allowing for moderate distinction in pollinator and plant traits. Our parameter set up best allows us to test the ability of phenological mechanisms to drive plasticity in pollinator specialization without a priori constraints on pollinator behavior though underlying baseline model parameters.

Baseline model sensitivity analysis has been thoroughly investigated by past studies[9,10,31] allowing us to focus analysis on effects of phenology. Parameter sweeps across phenological parameters were done with 37 combinations of bloom period and bloom break values for plant species. This created a distribution of temporal overlap in floral resources across plant species (see Metrics) that was then tested on bee phenologies composed of 56 combinations of flight period and flight break value. Each simulation of the 2072 possible combinations of the plant and animal phenologies was replicated 10 times using varied baseline (non-phenological) parameters (see paragraph above). While baseline model parameters vary across both plant and pollinator species within single simulations, all flowering and flight phenology parameters are shared across all plant and pollinator species respectively. This allows us to test the phenological mechanism against numerous baseline parameters per pollination interaction per simulation. Across all 2072 unique phenology parameter combinations simulated in each of our 3 network frameworks, each replicated 10 times, we simulated 62160 plant-pollinator networks representing over 22 million distinct pollination interactions.

*Choice of model bee genera & regions for empirical study*

*Andrena* (Hymenoptera: Andrenidae) and *Lasioglossum* (Hymenoptera: Halictidae) were selected as two contrasting model bee genera for this analysis. These two genera are the largest bee genera in the world[33], and both are well represented in Northern Hemisphere bee communities. *Andrena* contains around 1,500 species[34] and is distributed principally across the Holarctic with only a small number of species extending south to the Neotropics, sub-Saharan Africa and south-east Asia. *Lasioglossum* is a larger genus, containing nearly 1,800 species[35], and has a more cosmopolitan distribution being found in every continent except for Antarctica. The use of these large, widely distributed genera allows for comparisons across different biogeographical regions within two distinct bee lineages, each with a shared evolutionary history amongst its constituent species.

*Andrena* and *Lasioglossum* have notably different life history strategies. *Andrena* are protandrous, with males emerging in advance of the females but at the same time of the year. Females mate and then build nests and collect pollen and provision their offspring before dying. The pollen foraging period is short, typically lasing only a few weeks[36,37]. In contrast, already mated *Lasioglossum* females emerge in the spring and found nests. They collect pollen and rear offspring that often emerge and overlap with their activity period. Some species are eusocial, and will produce a brood of workers before producing reproductives[38]. *Lasioglossum* species display a range of social behaviors from solitary to eusocial[39-41], sometimes even within the same species across different parts of its range[42]. However, importantly, both social and solitary species of *Lasioglossum* have longer foraging periods than *Andrena*, typically spanning several months[36,37,42]. *Andrena* can consequently be considered a genus comprised of species with a 'short' foraging period, and *Lasioglossum* can be considered a genus of species with a 'long' foraging period, relative to one another (see also *Selection and Standardization of pollen load data*).

Both genera are found within the major clade of short-tongued bees within Apoidea[35]. Though many species of short-tongued bees from across this clade have independently evolved elongate proboscides to access nectar[43], morphological adaptations in *Andrena* or *Lasioglossum* bees for harvesting pollen are almost unknown. *Andrena* (*Scoliandrena*) and *Andrena* (*Hamandrena*) possess hooked hairs on the galea to pull pollen from the anthers of *Cryptantha* (Boraginaceae) and *Anchusa* (Boraginaceae) respectively, but these members of these subgenera are restricted to Western North America[44] and southern and eastern Europe[34,45] outside of our selected study regions of Britain and Michigan. In *Lasioglossum*, members of the subgenus *Sphecodogastra s.s.* are specialists on Onagraceae, and possess a scopa comprised of unbranched and sparse hairs and bristles rather than the typical scopa of branched or dense hairs[39]. These species are restricted to North America[39]. Although some species of *Andrena* and *Lasioglossum* preferentially forage from more morphologically complex flowers (e.g. Fabaceae, *Andrena wilkella* and *Lasioglossum lativentre*, ref 36), both genera as a whole favor shallow, open, and radially symmetrical flowers[36,43,46].

Plant species can have pollen that is difficult to digest by non-specialized bees[47], leading to patterns of specialization or avoidance[48]. Some plant species protect pollen physically by hiding it in deep tubes (Boraginaceae), or in poricidal anthers (Ericaceae, Solanaceae), or nototribic anthers (Lamiaceae) that deposit pollen on the bodies of bees[49]. There is therefore considerable variation in the rewards offered by different plant species. Conducting the analysis in Britain and Michigan, both of which are found in the Holarctic biogeographic region, means that whilst the constituent species of the respective floras differ, the botanical families present are very similar[50,51] and therefore likely to present rewards of a similar composition and availability. Furthermore, the two regions provide suitable extremes in their temporal resource overlap, making them ideal for addressing our hypothesis. The highly seasonal continental climate of Michigan[20] produces lower degrees of overlap and highly seasonal flowering communities, while the more mild oceanic climate of the British Isles[20] and its consistent temperatures produces longer, overlapping flowering times among plant species[21].

Restricting the analysis to the same genera across two regions also controls for possible impacts of differing travel costs when gathering resources. In broad terms, bee species foraging range is influenced by body size, with larger species able to travel greater distances to forage[52]. As *Andrena* and *Lasioglossum* communities contain species of the same size across both regions, potentially interacting effects of travel costs are minimized.

The use of both *Andrena* and *Lasioglossum* within two Holarctic regions therefore controls as much as possible for the possible effects of phylogenetic structuring that may affect the cost of travel, constraints of morphology, and unequal rewards between plant families. By comparing within and between these genera across the two regions of Britain and Michigan, the relative importance of foraging period and environmental seasonality in structuring a generalized foraging response in wild bees can be determined.

*Selection and standardization of pollen load data*

*Andrena* are characterized as short-season bees, collecting pollen over only a period of a few weeks. However, some species of *Andrena* express bivoltine behavior, having two reproductive generations in a single year. Although not unknown[53], bivoltinism is extremely rare in Nearctic *Andrena* and is not displayed by any of *Andrena* species in Michigan[54]. In contrast, bivoltinism is displayed by 13 (13/60, 22%) of extant British Andrena species and nine (9/29, 31%) of species included in our British dataset[37]. In order to prevent bivoltinism affecting our categorization of *Andrena* as bees with a short pollen foraging period, British *Andrena* with multiple generations were analyzed with each generation considered a separate species.

Data on British *Andrena* and *Lasioglossum* pollen collection come from the datasets compiled in Wood *et al.*[55] and Wood and Roberts[56]. Data on Michigan *Andrena* come from Wood and Roberts[46]. These were combined with new data on Michigan *Lasioglossum*, and additional new pollen load data for British *Lasioglossum*. Pollen was removed from these specimens and identified using light microscopy following the same method across all specimens following the methodology of Wood and Roberts[46]. The size of pollen loads on individual bees was estimated, ranging from a full load to a one-eighth load. Pollen grains were removed from the scopa using an entomological pin and transferred to a drop of water on a microscope slide. Grains were left to absorb water for a few minutes and then the slides were gently heated to allow evaporation. Molten glycerine jelly stained with fuchsin was added, and the slide was sealed with a coverslip. Following Müller and Kuhlmann[48], the percentage of the load composed of different plant species was estimated along three randomly selected lines across the cover slip at a magnification of ×400. The percentage of the load was estimated by the relative area of the slide occupied by each plant species, rather than the absolute number of grains[57]. Pollen species representing < 2% of the load were excluded from further analysis because their presence might have arisen from contamination. The percentages of pollen collected were corrected according to the overall size of each load to give a final weighting. Pollen loads were identified to the lowest taxonomic level possible using a reference collection assembled during the project, in most cases to genus.

Though pollen was identified to genus, the analyses were conducted at the botanical family level. Specialist bees, or oligoleges, collect all their pollen from a single botanical family[48], and this is the level at which most dietary specialization occurs in bees[8]. A minimum sample size of five pollen loads per species was selected as the cut-off for inclusion in this study in order to maximize the number of species included in the analysis. This sample size restriction meant that the first generation data for four British bivoltine *Andrena* was excluded, resulting in an effective sample size of 34 species in the British *Andrena* analysis (20 univoltine species, 5 species with data from both the spring and the summer generations, 4 species with data from the summer generation only).

All together, these criteria produced a final dataset of 2,561 pollen loads from 120 species, specifically 671 pollen loads from 51 Michigan *Andrena* species (63% of the *Andrena* fauna, ref 54), 355 pollen loads from 24 Michigan *Lasioglossum* species (30% of the *Lasioglossum* fauna), 815 pollen loads from 29 British Andrena species (49% of the *Andrena* fauna, ref 37), and 731 pollen loads from 16 British *Lasioglossum* species (50% of the *Lasioglossum* fauna).

Only one species was shared between the British and Michigan datasets, the Holarctic *Andrena barbilabris*. Because the British *Andrena* dataset compiled by Wood and Roberts[56] focused on polylectic species (species collecting pollen from multiple plant families), only two of the 29 species are oligoleges (2/29, 7%), a significantly smaller proportion than the extant British fauna (20/59, 34%, Chi squared, $\chi 2$=7.3, p=0.007). Following Wood and Roberts[46], 10 species of oligolectic *Andrena* were added to this dataset to bring this proportion in to line with the whole fauna (12/39, 31%, $\chi 2$=0.1, p=0.790). Because these are well-characterized oligolectic bees[36], no pollen load analysis was conducted for these species. There were no differences in the proportion of specialists in the *Lasioglossum* datasets and the entire regional fauna for Michigan ($\chi 2$=0.3, p=0.790) or Britain ($\chi 2$=0.0, p=1.000). Full details on the overall *Andrena* and *Lasioglossum* faunas of Britain and Michigan, their dietary status, and species with pollen load sample sizes included in this study can be found in Tables S6, S7, and S8.

*Metrics*

In model simulations, temporal resource overlap between any two plant species is defined as the shared area under each $R_i(t)$ curve (Fig S6a). The shared area can be found by:

1. Taking the minimum of any two $R_1$ and $R_2$.
2. This will give you a new curve, $\min(R_1, R_2)$ .
3. The area underneath this new curve $\min(R_1, R_2)$ is the resource overlap between $R_1$ and $R_2$ (shaded region in Fig S6a).
4. This resource overlap can be quantified by taking the integral of $\min(R_1, R_2)$. Integrals were taken using the trapezoidal method with unit spacing through the trapz function in Matlab.

Overlap per simulation was quantified as 1) total resource overlap summed across all $p_i$ (labeled $TRO$) and 2) average resource overlap experienced by each individual $p_i$ (labeled $ARO$). Simply counting overlapping shared time steps where $R_i > 0$ for two $R_i$ would be an insufficient metric because it would miss the quantity of resources that overlap in time. Our intuitive metric algorithm accounts for both shared time and quantity of overlapping resources. Modifying the bloom period ($b_p$) of plants with different bloom break values $b_{bv}$ provided a suitably even coverage of overlap values, whether measured as total overlap (Fig S6b) or average individual plant overlap. The exact overlap of a simulation at any given value of log(bs/bbv+1) will vary depending on the pollinator community.

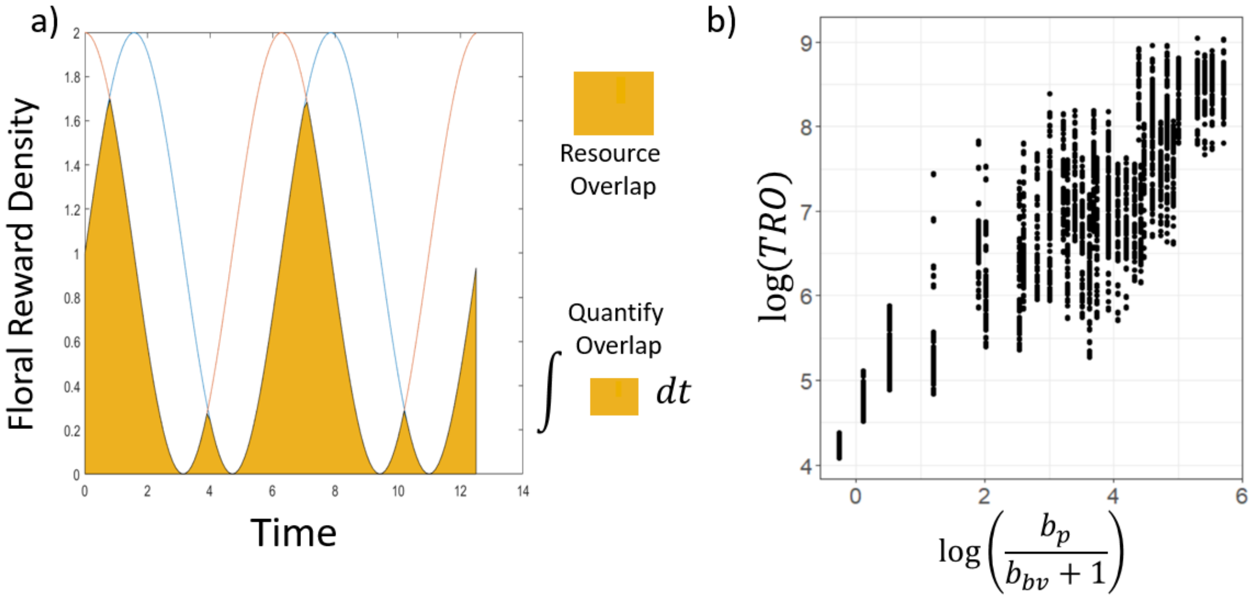


**Figure S6: Details of overlap metrics.** a) Diagram of overlap defined by two $R_i(t)$ curves. Two example plant species rewards production curves across time ($R_i(t)$ are shown in red and blue. Shared area under the curve is considered the overlapping resources and is shaded orange. Quantifying this area is done by taking the integral of the shared area. b) Example from 58$p$ x 100$a$ web simulations of range in total resource overlap ($TRO$) provided by separating bloom period ($b_p$) values by bloom break values ($b_{bv}$). The denominator adds 1 to $b_{bv}$ avoid dividing by 0 when $b_{bv} = 0$. Each dot represents overlap per simulation at different $b_p$ and $b_{bv}$ values. The range in overlap per combination of $b_p$ and $b_{bv}$ comes from the different phenology metrics tested for the pollinators.

Defining the confines of a specialist or generalist has been a long-running debate in ecology and evolutionary biology, with different sub-fields often using different metrics. For example, phylogenetic studies have found strong evidence of herbivorous insects specializing on plants at the family level[13], while in other fields, such as network ecology, specialists and generalists have been defined by the number of species interactions through the degree distribution of links in the network[58]. In order to consider the range of ideas used in defining specialists and generalists, we have employed multiple metrics to measured specialization that can be applied to both the foraging effort ($\alpha$) based metrics of the model and the pollen loads from our empirical data.

We label the first metric as "Deviation from Generalism" or DFG. For a single pollinator species, DFG is defined as the degree away from completely even foraging efforts or pollen loads across all possible resources. Specifically, DFG for a pollinator species $j$ is defined as the sum of pairwise differences of foraging effort across all plant species divided the number of plant species minus 1. Parametrically, we can write:

$$DFG_{a_j} = \frac{\sum_{i,k\epsilon p}|\alpha_{ij}-\alpha_{kj}|}{N_p-1} \qquad \text{Eq (10)}$$

where DFG ranges from [0,1] with a perfect generalist scoring 0 and a complete specialist scoring 1. An example is provided in Table S5.

**Table S5:** Example foraging preference matrix used to calculate DFG for pollinators from simulations (using $\alpha$) or pollen load data. In the case of empirical pollen load data, the percent of pollen load is either grouped by plant family or plant genus.

| **Pollinators (model)** | **Effort on Plant 1** | **Effort on Plant 2** | **Effort on Plant 3** | **Effort on Plant 4** | **DFG** |
|---|---|---|---|---|---|
| **Pollinators (pollen)** | **Plant 1 Pollen %** | **Plant 2 Pollen %** | **Plant 3 Pollen %** | **Plant 4 Pollen %** | **DFG** |
| Bee sp. 1 | 1.0 | 0 | 0 | 0 | 1 |
| Bee sp. 2 | 0.5 | 0.5 | 0 | 0 | 0.667 |
| Bee sp. 3 | 0.2 | 0.4 | 0.2 | 0.2 | 0.2 |
| Bee sp. 4 | 0.25 | 0.25 | 0.25 | 0.25 | 0 |

In this example, the DFG scores would be calculated as follows:

$$DFG_1 = \frac{|1-0|+|1-0|+|1-0|+|0-0|+|0-0|+|0+0|}{3} = \frac{3}{3} = 1$$

$$DFG_2 = \frac{|0.5-0.5|+|0.5-0|+|0.5-0|+|0.5-0|+|0.5-0|+|0-0|}{3} = \frac{2}{3} = 0.667$$

$$DFG_3 = \frac{|0.2-0.4|+|0.2-0.2|+|0.2-0.2|+|0.4-0.2|+|0.4-0.2|+|0.2-0.2|}{3} = \frac{0.6}{3} = 0.2$$

$$DFG_4 = \frac{|0.25-0.25|+|0.25-0.25|+|0.25-0.25|+|0.25-0.25|+|0.25-0.25|+|0.25-0.25|}{3} = \frac{0}{3} = 0$$

When the DFG metric is used with pollen load data, the pollen amounts from each plant group in a bee species' overall pollen load is broken into percentages and substituted for the model's foraging effort (Table S5), under the assumption that higher pollen loads of any particular plant group generally correspond to higher foraging effort on that group. The term group is used here because pollen data was evaluated against both family and genus level plant data. The algorithm used to calculate our DFG score is a holistic accounting for every possible interaction and compares all possible interactions amongst each other to accumulate a score. However, it may not be the best option in all circumstances. While not necessarily requiring a complete accounting of all available resources, DFG's [0,1] range and attempt to calculate specialization while considering all potential foraging options means DFG functions best when available data can convincingly represent all possible foraging options. This is not necessarily guaranteed in empirical surveys.

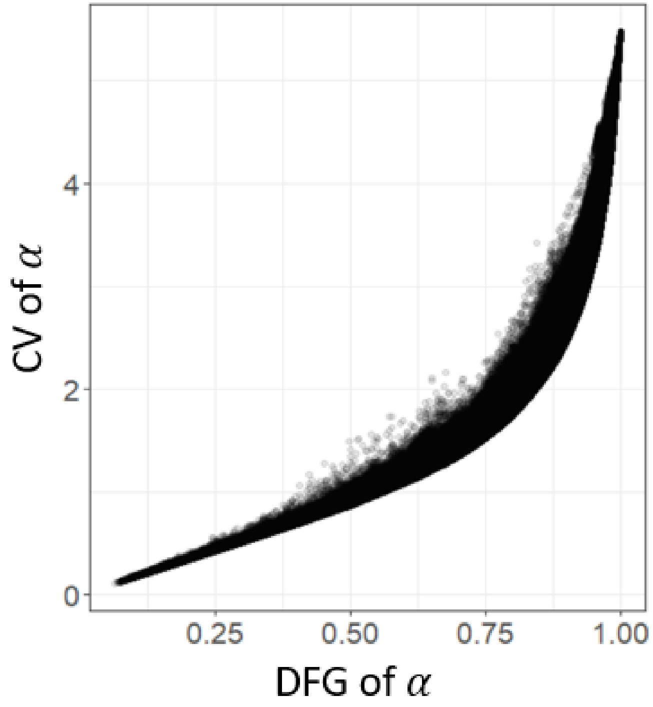


**Figure S7: Comparison of specialization metrics.** Comparison of DFG and CV from all model simulations in the 2nd largest network tested. Each dot represents a measurement from a single pollinator taken at the end of the simulation which in this case represents 7,061,376 data points.

In light of the above, we also employed the coefficient of variance (CV) as our second metric of specialization. With model simulations, we consider the CV of pollinators' $\alpha$ values whereas in the empirical data we take CV of the percent of pollen load across plant groups. As a metric of specialization, the CV will increase as when there are a limited number of extreme values. It performs the same whether or not the full range of possible resource options for foraging pollinators is known. The relationship between the two metrics is positive monotonic but non-linear (Fig S7) showing that the two metrics relate to the measurement of specialization differently as desired.

In measuring model output, both DFG and CV metrics are measured in two ways. First each metric is taken on the $\alpha$ values at the end of each simulation ($DFG_{End}$ & $CV_{End}$ taken at t=6000). Second, each metric is run on the $\alpha$ values averaged across the last 1000 time steps of each simulation ($DFG_{Avg}$ & $CV_{Avg}$). This is done to provide multiple points of comparison to our empirical pollen load data. One could argue that pollen loads taken over multiple years would be better compared to model data taken across an extended period of time steps. On the other hand, it could also be argued that foraging bees collected for data are exhibiting an innate search pattern that is the end result of generations of evolved behavior, better compared with the end state of simulations. Either way, the two methods are intuitively correlated and using either supports our qualitative conclusions.

In measuring empirical pollen load data, we measured DFG and CV of bee species by grouping plants by either family level or genus level. Network studies in pollination ecology, including our model, tend to group plants at the species level. While, identifying pollen to species level is prohibitively difficult, genus level identifications allow for a closer approximation. However, studies focused on specialization in herbivores (particularly insects) have centered around grouping plants at the family level[13,48,57]. Given the range, we decided to use both taxonomic levels and found that results were qualitatively similar regardless of grouping.

*Statistics*

Statistical analysis was done using R version 3.3.2[59]. Differences in CV and DFG scores between bee genera and geographic regions were tested using Kruskal-Wallis one-way analysis of variance tests.

Because both the CV and DFG scores were calculated from deeply non-normal underlying data (necessarily so since patterns of diet breadth are decidedly non-normal at the botanical family level, Forister *et al*. ref 13), it was not possible to transform them sufficiently to allow for parametric statistical assessment. Regression analysis on model diet breadth results was done using the lm function and measured with traditional $R^2$ metric when working with $DFG_{End}$ results or with beta regressions (betareg package in R) and the pseudo $R^2$ metric when working with $DFG_{Avg}$. When measuring diet breadth results with $CV_{End}$ or $CV_{Avg}$, log-linked gamma regressions were completed with the mgcv package and measured with the for the amount of deviance accounted for by the regression model with the $D^2$ metric[60].

*Code availability & simulation information*

Simulation code, metric code, and simulation data are available at the repository https://github.com/fsvaldovinos/Phenology.

**Supplementary Discussion**

When considering the factors that drive specialization in insect herbivores, it is impossible not to consider the role of plant chemistry. Phytochemical diversity, the diversity of plant compounds including anti-herbivory defense compounds, has been strongly associated with both insect herbivore diversity and dietary specialization[19,22,23]. For many insect herbivores, their diversity is much greater in tropical environments. This is particularly well studied in Lepidoptera where both their species richness and degree of specialization increases from temperate to tropical regions[22,24]. However, a completely different pattern is seen in the bees where tropical diversity and specialization is mediocre compared to diversity in Mediterranean and xeric environments where it is high[8,25,26]. Indeed, tropical environments are dominated by highly generalized and often social species that visit a huge variety of botanical families (e.g. ref 27, 28, 61).

Though the theoretical impact of variable quality between different resources is clear[7,8], bees have an ecology that fundamentally differs from most insect herbivores. They are nest makers, and they collect pollen as adults before feeding it to their offspring. Therefore, even though their larvae are immobile, they are effectively mobile because they have the potential to feed on pollen collected from many botanical families in a way that most larvae which must feed themselves are not, due to their limited mobility during this stage of their development. In this regard bee larvae are functionally closer to those insects that are herbivorous during their adult stages such as Chrysomelid beetles[62], and indeed these groups show higher levels of generalization[13].

The ability to feed their offspring from multiple combined sources has led to the suggestion that bees may adaptively pollen mix to negate the harmful properties of a particular pollen type that would be lethal if consumed as the sole source of larval nutrition[63]. This mixing of different resources is however ultimately also affected by resource overlap, as it does not matter how much mobility an organism has if there are still limited different resources available to it during the course of its lifespan. Indeed, pollen mixing of botanical families identified as having negative pollen properties as a behavior is found in the *Andrena* of the UK but not those in Michigan[46]. This is not to say that plant chemistry is unimportant for bees, and indeed there are many examples from within this group of plant chemistry driving and indeed restricting host plant choice[47,64]. Instead, we argue that the fundamental difference in the life stage at which herbivory occurs means that for bees, inherent resource quality is relatively less important than resource availability, as defined by the balance between their inherent flight period and local flowering patterns. Clearly, despite existing in the same extremely phytochemically diverse landscapes, tropical bees have not evolved anything approaching the same high degree of dietary specialization as seen in the tropical Lepidoptera.

This weaker relationship in dietary specialization in bees compared to other insect herbivores is illustrated by directly comparing the overall pattern of specialization at the botanical family level. Species in Michigan adhere more closely to a power law relationship, see Supplementary Figure S11a,c, similar to Forister et al[13], but the pattern is much looser and closer to a zero-inflated Poisson distribution (Supplementary Figure S11b, d). Constraining the number of dietary choices available to a mobile species within its lifespan by reducing flowering overlap is directly analogous to reducing its mobility. Even if it can fly, if there is only one flowering plant available then it is functionally the same as being an egg laid on a plant; their choices are limited. We would therefore expect that phytochemical diversity and other factors that influence resource quality would show the greatest impact on bee diets in environments where their choices are limited by flowering patterns. More studies on patterns of pollen collection in bees from a wider variety of global environments are needed, but it would seem that the biggest driver of the divergent patterns of herbivory seen in bees is their highly mobile period of pollen foraging as adults.

### Supplementary Figures and Tables

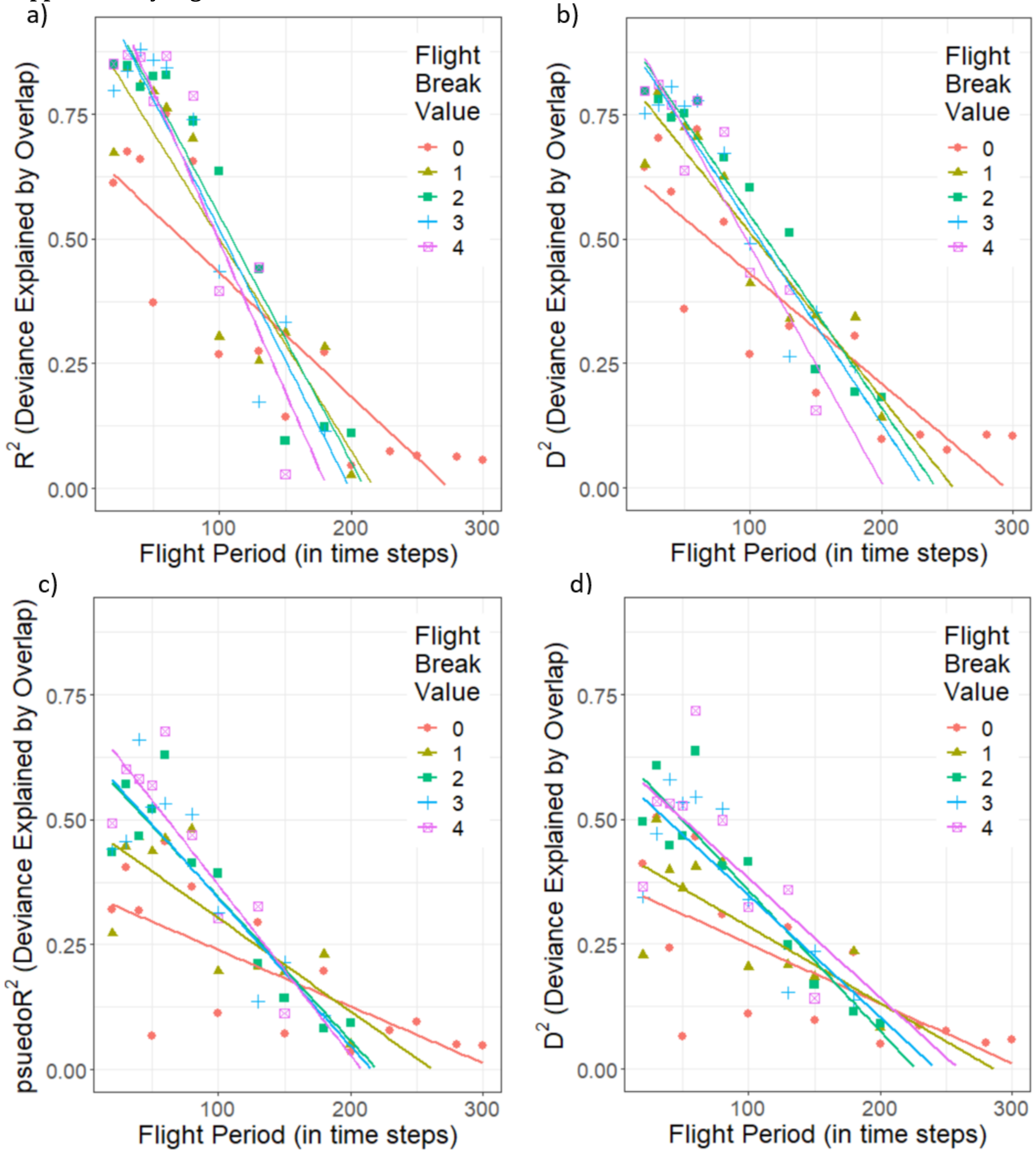


**Figure S8: Results summary in $N_p = 30$ plants & $N_a = 50$ framework.** Diet breadth deviance explained by average resource overlap ($ARO$) across flight periods ($f_p$) and flight break values ($f_{bv}$). a) The $R^2$ values of diet breadth measured through $DFG_{End}$regressed against $ARO$ in linear models across different flight period ($f_p$) and flight break values ($f_{bv}$). Regression lines plotted for visual aid. b) The $D^2$ values of diet breadth measured through $CV_{End}$ regressed against $ARO$ in generalized linear models (Gamma distribution w/ log link) across different flight period ($f_p$) and flight break vales ($f_{bv}$). Regression lines plotted for visual aid. c) The pseudo $R^2$ values of diet breadth measured through

$DFG_{Avg}$ regressed against $ARO$ in beta regression models across different flight period ($f_p$) and flight break values ($f_{bv}$). Regression lines plotted for visual aid. b) The $D^2$ values of diet breadth measured through $CV_{Avg}$ regressed against $ARO$ in generalized linear models (Gamma distribution w/ log link) across different flight period ($f_p$) and flight break vales ($f_{bv}$). Regression lines plotted for visual aid. Overall, regardless of $f_{bv}$, there is a decrease in the effect of temporal resource overlap on diet breadth as the $f_s$ of pollinators increases. Data taken from network framework w/ $N_p = 30$ plants & $N_a = 50$.

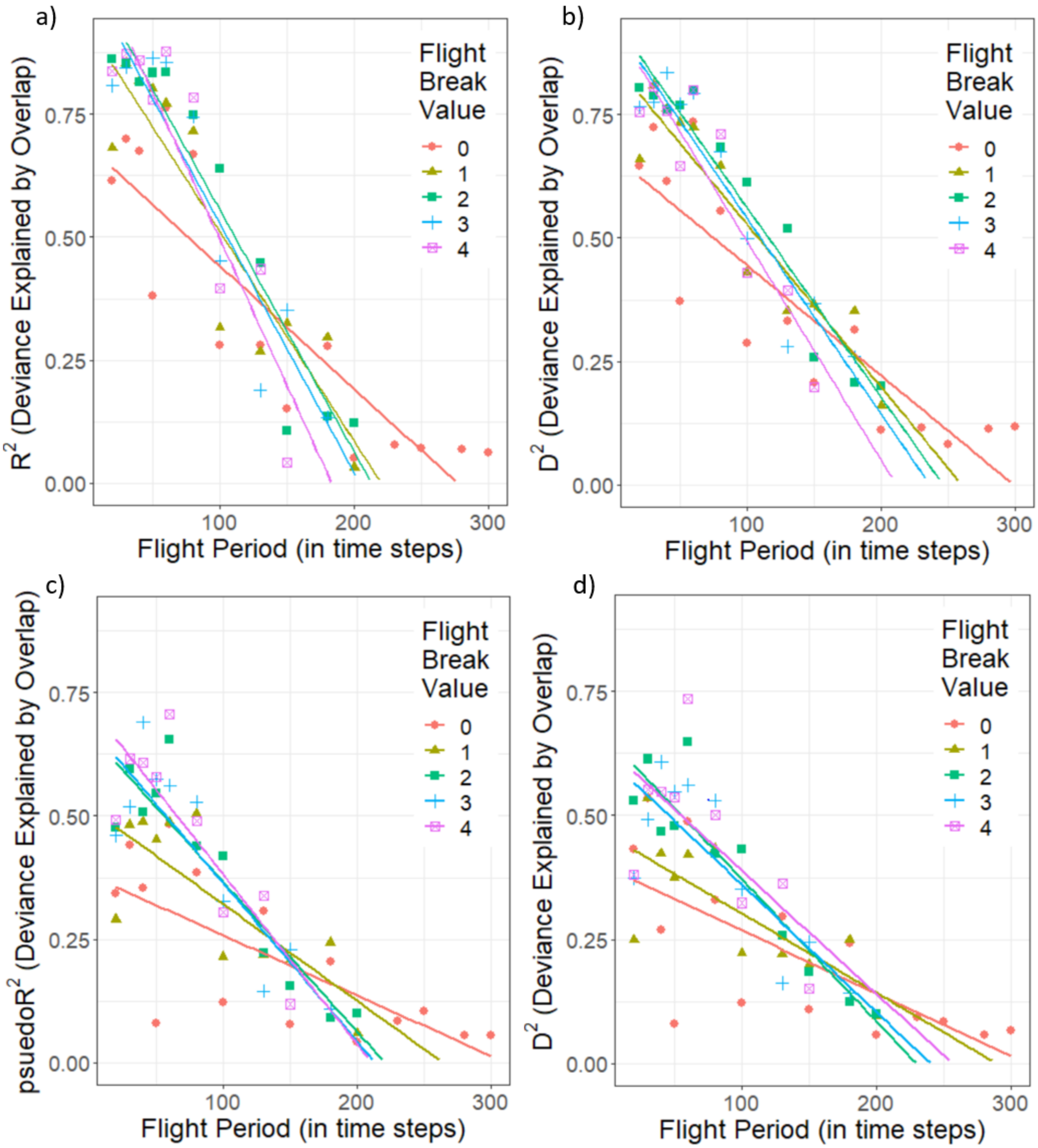


**Figure S9: Results summary in $N_p = 48$ plants & $N_a = 71$ framework.** Diet breadth deviance explained by average resource overlap ($ARO$) across flight periods ($f_p$) and flight break values ($f_{bv}$). a) The $R^2$ values of diet breadth measured through $DFG_{End}$regressed against $ARO$ in linear models across different flight period ($f_p$) and flight break values ($f_{bv}$). Regression lines plotted for visual aid. b) The $D^2$ values of diet breadth measured through $CV_{End}$ regressed against $ARO$ in generalized linear models (Gamma distribution w/ log link) across different flight period ($f_p$) and flight break vales ($f_{bv}$). Regression lines plotted for visual aid. c) The pseudo $R^2$ values of diet breadth measured through

$DFG_{Avg}$ regressed against $ARO$ in beta regression models across different flight period ($f_p$) and flight break values ($f_{bv}$). Regression lines plotted for visual aid. b) The $D^2$ values of diet breadth measured through $CV_{Avg}$ regressed against $ARO$ in generalized linear models (Gamma distribution w/ log link) across different flight period ($f_p$) and flight break vales ($f_{bv}$). Regression lines plotted for visual aid. Overall, regardless of $f_{bv}$, there is a decrease in the effect of temporal resource overlap on diet breadth as the $f_s$ of pollinators increases. Data taken from network framework w/ $N_p = 48$ plants & $N_a = 71$.

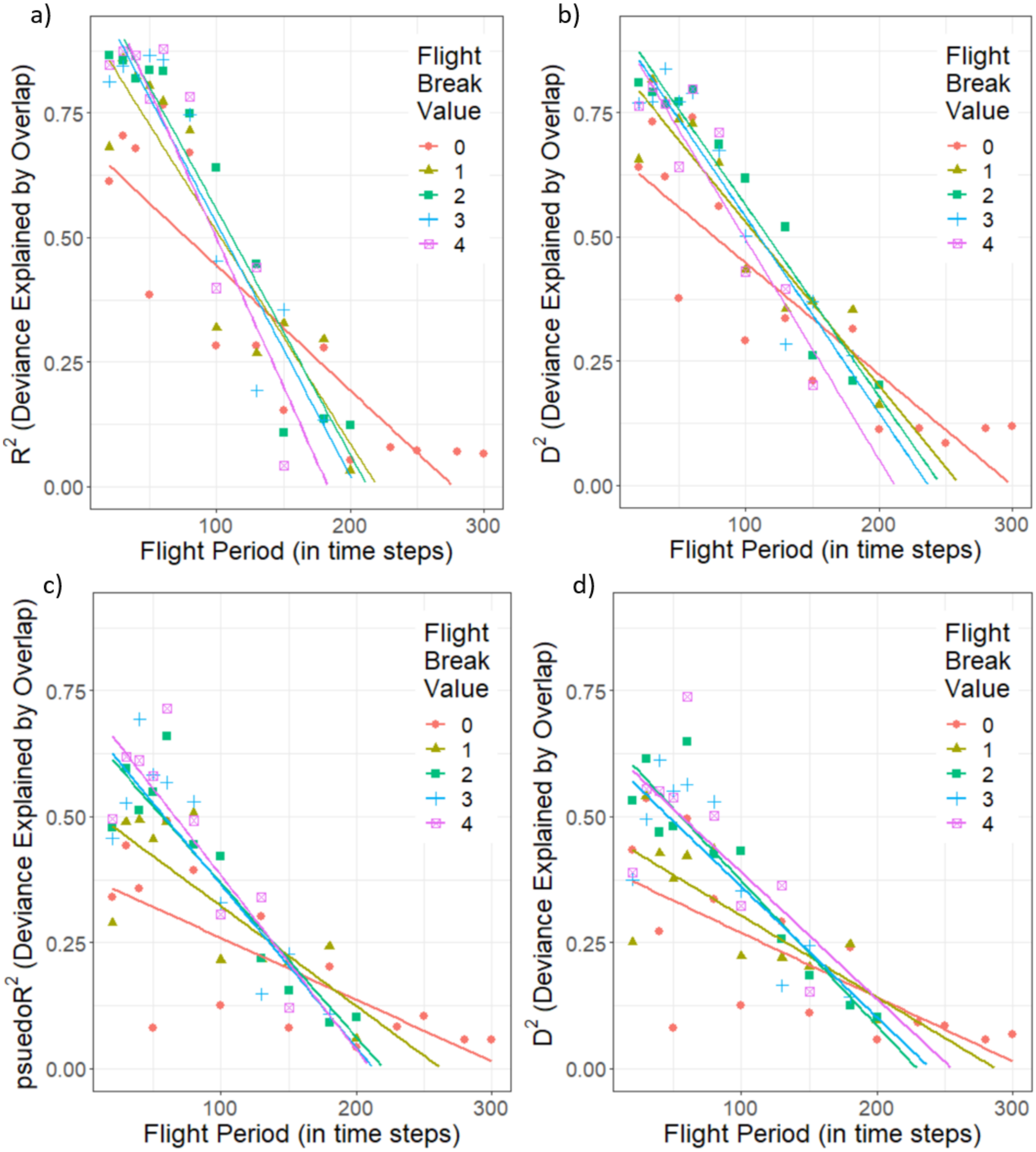


**Figure S10: Results summary in $N_p = 58$ plants & $N_a = 100$ framework.** Diet breadth deviance explained by average resource overlap (*ARO*) across flight periods ($f_p$) and flight break values ($f_{bv}$). a) The $R^2$ values of diet breadth measured through $DFG_{End}$ regressed against *ARO* in linear models across different flight period ($f_p$) and flight break values ($f_{bv}$). Regression lines plotted for visual aid. b) The $D^2$ values of diet breadth measured through $CV_{End}$ regressed against *ARO* in generalized linear models (Gamma distribution w/ log link) across different flight period ($f_p$) and flight break vales ($f_{bv}$). Regression lines plotted for visual aid. c) The pseudo $R^2$ values of diet breadth measured through $DFG_{Avg}$ regressed against *ARO* in beta regression models across different flight period ($f_p$) and flight break values ($f_{bv}$). Regression lines plotted for visual aid. b) The $D^2$ values of diet breadth measured through $CV_{Avg}$ regressed against $ARO$ in generalized linear models (Gamma distribution w/ log link) across different flight period ($f_p$) and flight break vales ($f_{bv}$). Regression lines plotted for visual aid. Overall, regardless of $f_{bv}$, there is a decrease in the effect of temporal resource overlap on diet breadth as the $f_s$ of pollinators increases. Data taken from network framework w/ $N_p = 58$ plants & $N_a = 100$.

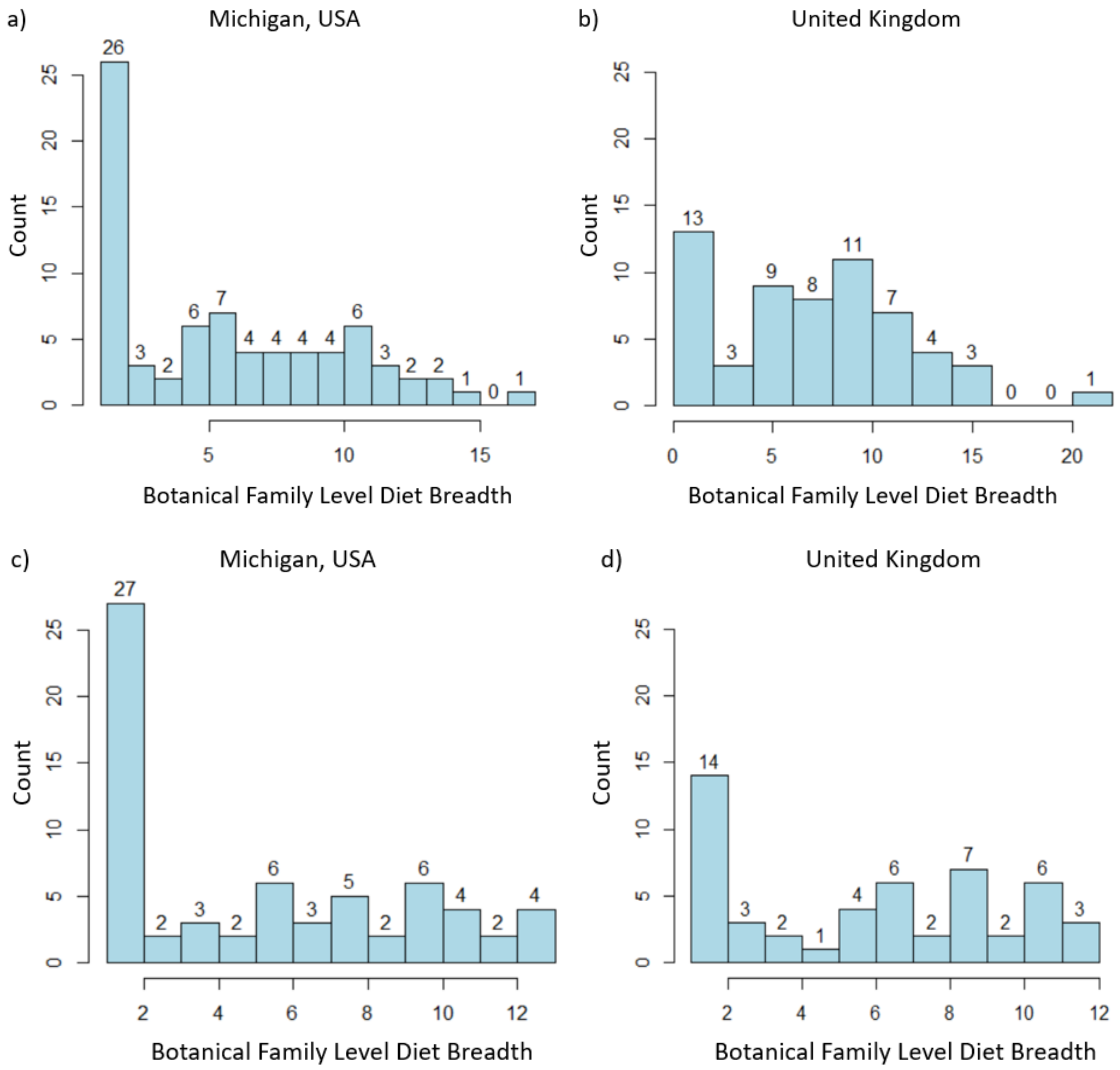


**Figure S11: Diet breadth distributions.** Botanical family level diet breadth for studied bee species following the format of Forister *et al*[13]. Data are from a) raw Michigan, USA data, b) raw United Kingdom data, c) rarefied Michigan, USA data, d) rarefied United Kingdom data. Data rarefaction followed Wood and Roberts[56], and samples were rarefied to the minimum sample size of 10 pollen loads. Bivoltine *Andrena* generations were treated as separate species. This resulted in a dataset of 2,524 pollen loads from 113 species (functionally 116 species).

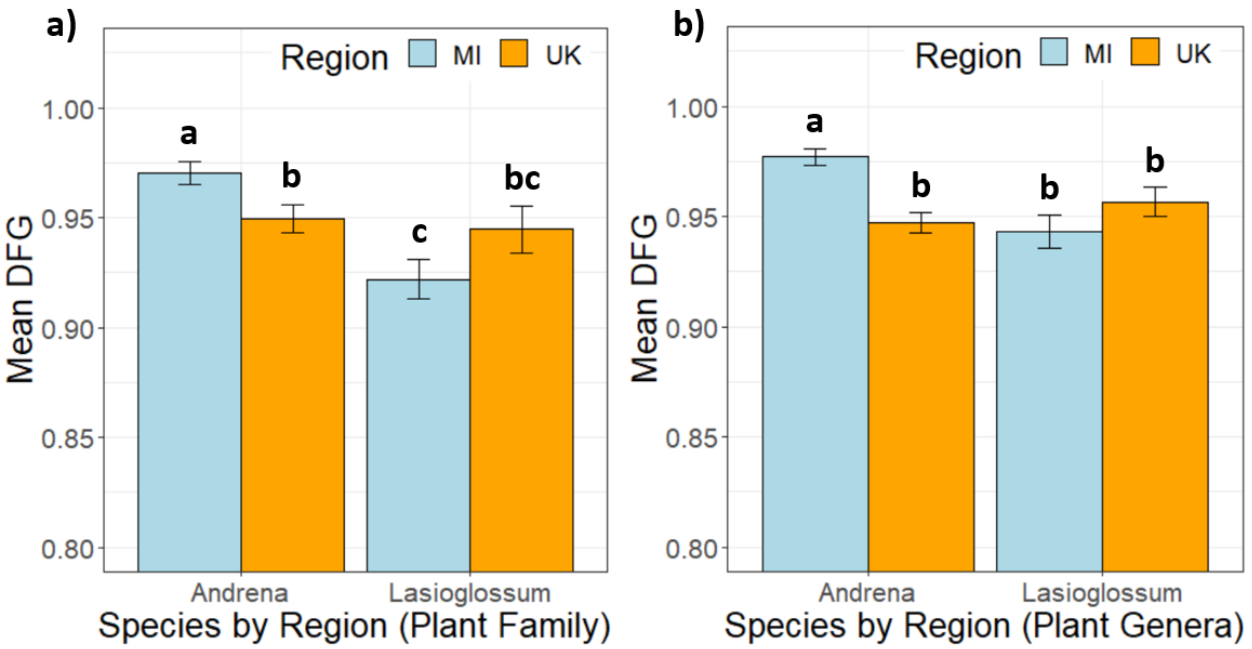


**Figure S12: Specialization across regions using Departure from Generalism (DFG).** Difference in dietary specialization of *Andrena* and *Lasioglossum* bees between Michigan and the UK at (a) the botanical family and (b) botanical genus level, with higher DFG score indicating greater average specialization per bee species. Bars headed by different letters are significantly different ($p<0.05$) error bars show standard error.

**Table S6.** Extant *Andrena* species of Michigan and Britain and their dietary status. Species are ordered by subgenus. Species without a listed host range are polylectic.

| Subgenus | Michigan species | Host range | British species | Host range |
|---|---|---|---|---|
| *Andrena s. str.* | *carolina* Viereck | *Vaccinium* | *apicata* Smith | *Salix* |
| | *clarkella* Kirby | *Salix* | *clarkella* Kirby | *Salix* |
| | *frigida* Smith | *Salix* | *fucata* Smith | |
| | *mandibularis* Robertson | | *fulva* Müller | |
| | *milwaukeensis* Graenicher | | *helvola* L. | |
| | *rufosignata* Cockerell | | *lapponica* Zetterstedt | |
| | *thaspii* Graenicher | | *praecox* Scopoli | *Salix* |
| | *tridens* Robertson | | *synadelpha* Perkins | |
| | | | *varians* Rossi | |
| *Callandrena s.l.* | *aliciae* Robertson | Asteraceae | | |
| | *asteris* Robertson | *Solidago* | | |
| | *gardineri* Cockerell | *Packera* | | |
| | *helianthi* Robertson | *Helianthus* | | |
| | *krigiana* Robertson | *Krigia* | | |
| | *placata* Mitchell | *Solidago* | | |
| | *rudbeckiae* Robertson | *Rudbeckia* | | |
| | *simplex* Smith | *Solidago* | | |
| *Charitandrena* | | | *hattorfiana* Fabricius | Dipsacaceae |
| *Chlorandrena* | | | *humilis* Imhoff | Asteraceae |
| *Chrysandrena* | | | *fulvago* Christ | Asteraceae |
| *Cnemidandrena* | *canadensis* Dalla Torre | *Solidago* | *denticulata* Kirby | Asteraceae |
| | *chromotricha* Cockerell | Asteraceae | *fuscipes* Kirby | Ericaceae |
| | *hirticincta* Provancher | *Solidago* | *nigriceps* Kirby | |
| | *nubecula* Smith | *Solidago* | *simillima* Smith | |
| | *parnassiae* Cockerell | *Parnassia* | | |
| | *peckhami* Cockerell | *Helianthus* | | |
| | *robervalensis* Mitchell | *Solidago* | | |
| *Conandrena* | *bradleyi* Viercek | *Vaccinium* | | |
| *Euandrena* | *algida* Smith | *Salix* | *bicolor* Fabricius | |
| | *geranii* Robertson | *Hydrophyllum* | *ruficrus* Nylander | *Salix* |
| | *nigrihirta* Ashmead | | | |
| | *polemonii* Robertson | *Polemonium* | | |
| *Gonandrena* | *fragilis* Smith | *Cornus* | | |
| | *integra* Smith | *Cornus* | | |
| | *persimulata* Viereck | *Cornus* | | |
| | *platyparia* Robertson | *Cornus* | | |
| *Holandrena* | *cressonii* Robertson | | *labialis* Kirby | |
| *Hoplandrena* | | | *bucephala* Stephens | |
| | | | *ferox* Smith | *Quercus* |
| | | | *rosae* Panzer | |
| | | | *scotica* Perkins | |
| | | | *trimmerana* Kirby | |
| *Iomelissa* | *violae* Robertson | *Viola* | | |
| *Larandrena* | *miserabilis* Cresson | | | |
| *Leucandrena* | *barbilabris* Kirby | | *argentata* Smith | |
| | *erythronii* Robertson | *Erythronium* | *barbilabris* Kirby | |
| *Margandrena* | | | *marginata* Fabricius | Dipsacaceae |
| *Melandrena* | *carlini* Cockerell | | *cineraria* L. | |
| | *commoda* Smith | | *nigroaenea* Kirby | |
| | *dunningi* Cockerell | | *nitida* Müller | |
| | *hilaris* Smith | | *thoracica* Fabricius | |
| | *nivalis* Smith | | *vaga* Panzer | *Salix* |

| | | | | |
|---|---|---|---|---|
| | *pruni* Robertson | | | |
| | *regularis* Malloch | | | |
| | *vicina* Smith | | | |
| *Micrandrena* | *illinoensis* Robertson | *Salix* | *alfkenella* Perkins | |
| | *melanochroa* Cockerell | | *falsifica* Perkins | |
| | *neonana* Viereck | | *minutula* Kirby | |
| | *nigrae* Robertson | | *minutuloides* Perkins | |
| | *personata* Robertson | | *niveata* Friese | Brassicaceae |
| | *salictaria* Robertson | *Salix* | *semilaevis* Pérez | |
| | *ziziae* Robertson | Apiaceae | *subopaca* Nylander | |
| *Notandrena* | | | *chrysosceles* Kirby | |
| | | | *nitidiuscula* Schenck | Apiaceae |
| *Oreomelissa* | | | *coitana* Kirby | |
| *Parandrena* | *andrenoides* Cresson | *Salix* | | |
| | *wellesleyana* Robertson | *Salix* | | |
| *Plastandrena* | *crataegi* Robertson | | *bimaculata* Kirby | |
| | | | *nigrospina* Thompson | Brassicaceae |
| | | | *pilipes* Fabricius | |
| | | | *tibialis* Kirby | |
| *Poecilandrena* | | | *labiata* Fabricius | |
| *Poliandrena* | | | *florea* Fabricius | *Bryonia* |
| *Proxiandrena* | | | *proxima* Kirby | Apiaceae |
| *Ptilandrena* | *distans* Provancher | *Geranium* | *angustior* Kirby | |
| | *erigeniae* Robertson | *Claytonia* | | |
| *Rhacandrena* | *brevipalpis* Cockerell | *Rhus* | | |
| | *robertsonii* Dalla Torre | | | |
| *Scaphandrena* | *arabis* Robertson | Brassicaceae | | |
| *Scrapteropsis* | *alleghaniensis* Viereck | | | |
| | *imitatrix* Cresson | | | |
| *Simandrena* | *nasonii* Robertson | | *congruens* Schmiedeknecht | |
| | *wheeleri* Graenicher | | *dorsata* Kirby | |
| *Taeniandrena* | | | *ovatula* Kirby | |
| | | | *similis* Smith | Fabaceae |
| | | | *wilkella* Kirby | Fabaceae |
| *Tarsandrena* | | | *tarsata* Nylander | *Potentilla* |
| *Thysandrena* | *bisalicis* Viereck | *Salix* | | |
| | *w-scripta* Viereck | | | |
| *Trachandrena* | *ceanothi* Viereck | | *haemorrhoa* Fabricius | |
| | *forbesii* Robertson | | | |
| | *heraclei* Robertson | | | |
| | *hippotes* Robertson | | | |
| | *mariae* Robertson | *Salix* | | |
| | *mirandra* Smith | | | |
| | *nuda* Robertson | | | |
| | *rehni* Viereck | | | |
| | *rugosa* Cockerell | | | |
| | *sigmundi* Cockerell | *Salix* | | |
| | *spiraeana* Robertson | | | |
| | *virginiana* Mitchell | | | |
| *Tylandrena* | *erythrogaster* Ashmead | *Salix* | | |
| | *perplexa* Smith | | | |
| | *wilmattiae* Cockerell | | | |
| *Zonandrena* | | | *flavipes* Panzer | |
| | | | *gravida* Imhoff | |
| Total: | 81 | 42 | 59 | 20 |
| Proportion oligolectic: | | 51.9 | | 33.9 |

**Table S6 Notes.** Wood and Roberts[46] added 11 oligolectic species to their analysis, including *Andrena lapponica* Zetterstedt. However, Müller[65] demonstrated that this species is polylectic, and so it was not used in the analysis presented in the current paper.

Gibbs *et al*.[54] listed *Andrena barbara* Bouseman & LaBerge, *A. morrisonella* Viereck, *A. confederata* Viereck and *A. wilkella* Kirby as part of the Michigan fauna. Examination of material at the University of Michigan Museum of Zoology (Ann Arbor, Michigan) has shown that the record of *A. barbara* is based on a misidentification (TJW, *unpublished data*). The status of *Andrena morrisonella* is still taxonomically uncertain and females cannot be reliably separated from *A. imitatrix*[66]. *Andrena confederata* was recorded from Michigan on the basis of a single male that cannot be found[54], and repeated searching at the locality has not produced any more records of this species. *Andrena wilkella* is introduced to North America[67]. These species have therefore been excluded from the Michigan fauna for the purposes of this study.

**Table S7.** Extant pollen-collecting *Lasioglossum* species of Michigan and Britain and their dietary status. Species are ordered by subgenus. Species without a listed host range are polylectic.

| Subgenus | Michigan species | Host range | British species | Host range |
|---|---|---|---|---|
| *Dialictus* | *abanci* Crawford | | *cupromicans* Pérez | |
| | *achilleae* Mitchell | | *leucopus* Kirby | |
| | *admirandum* Sandhouse | | *morio* Fabricius | |
| | *albipenne* Robertson | | *smeathmanellum* Kirby | |
| | *anomalum* Robertson | | | |
| | *atwoodi* Gibbs | | | |
| | *bruneri* Crawford | | | |
| | *callidum* Sandhouse | | | |
| | *cattellae* Ellis | | | |
| | *ceanothi* Mitchell | | | |
| | *coeruleum* Robertson | | | |
| | *coreopsis* Robertson | | | |
| | *cressonii* Robertson | | | |
| | *dreisbachi* Mitchell | | | |
| | *ellisae* Sandhouse | | | |
| | *ephialtum* Gibbs | | | |
| | *fattigi* Mitchell | | | |
| | *floridanum* Robertson | | | |
| | *foveolatum* Robertson | | | |
| | *gotham* Gibbs | | | |
| | *hartii* Robertson | | | |
| | *heterognathum* Mitchell | | | |
| | *hitchensi* Gibbs | | | |
| | *illinoense* Robertson | | | |
| | *imitatum* Smith | | | |
| | *laevissimum* Smith | | | |
| | *leucocomum* Lovell | | | |
| | *lineatulum* Crawford | | | |
| | *nigroviride* Graenicher | | | |
| | *novascotiae* Mitchell | | | |
| | *oblongum* Lovell | | | |
| | *obscurum* Robertson | | | |
| | *oceanicum* Cockerell | | | |
| | *paradmirandum* Knerer & Atwood | | | |
| | *perpunctatum* Ellis | | | |
| | *pictum* Crawford | | | |
| | *pilosum* Smith | | | |
| | *planatum* Lovell | | | |
| | *pruinosum* Robertson | | | |
| | *rufulipes* Cockerell | | | |
| | *sagax* Sandhouse | | | |
| | *sheffieldi* Gibbs | | | |
| | *smilacinae* Robertson | | | |
| | *subversans* Mitchell | | | |
| | *subviridatum* Cockerell | | | |
| | *taylorae* Gibbs | | | |
| | *tegulare* Robertson | | | |
| | *tenax* Sandhouse | | | |
| | *timothyi* Gibbs | | | |
| | *trigeminum* Gibbs | | | |
| | *versans* Lovell | | | |

| | | | | |
|---|---|---|---|---|
| | *versatum* Robertson | | | |
| | *vierecki* Crawford | | | |
| | *viridatum* Lovell | | | |
| | *weemsi* Mitchell | | | |
| | *zephyrum* Smith | | | |
| *Evylaeus* s.s. | *cinctipes* Provancher | | | |
| *Hemihalictus* | *birkmanni* Crawford | | *angusticeps* Perkins | |
| | *fedorense* Crawford | | *brevicorne* Schenck | Asteraceae |
| | *foxii* Robertson | | *minutissimum* Kirby | |
| | *inconditum* Cockerell | | *nitidiusculum* Kirby | |
| | *lustrans* Cockerell | Asteraceae | *parvulum* Schenck | |
| | *macoupinense* Robertson | | *pauperatum* Brullé | |
| | *nelumbonis* Robertson | Nymphaceae | *punctatissimum* Schenck | |
| | *pectorale* Smith | | *puncticolle* Morawitz | |
| | *swenki* Crawford | | *rufitarse* Zetterstedt | |
| | | | *semilucens* Alfken | |
| | | | *sexstrigatum* Kirby | |
| | | | *villosulum* Kirby | Asteraceae |
| *Lasioglossum s. str.* | *acuminatum* McGinley | | *laevigatum* Kirby | |
| | *athabascense* Sandhouse | | *lativentre* Schenck | |
| | *coriaceum* Smith | | *prasinum* Smith | |
| | *forbesii* Robertson | | *quadrinotatum* Kirby | |
| | *fuscipenne* Smith | | *xanthopus* Kirby | |
| | *paraforbesii* McGinley | | | |
| *Leuchalictus* | | | *leucozonium* Schrank | |
| | | | *zonulum* Smith | |
| *Sphecodogastra* | *comagenense* Knerer & Atwood | | *albipes* Fabricius | |
| | *oenothera* Stevens | *Oenothera* | *calceatum* Scopoli | |
| | *quebecense* Crawford | | *fratellum* Pérez | |
| | *seillean* Gibbs & Packer | | *fulvicorne* Kirby | |
| | *texanum* Cresson | *Oenothera* | *laticeps* Schenck | |
| | *truncatum* Robertson | | *malachurum* Kirby | |
| | | | *pauxillum* Schenck | |
| Total: | 77 | 4 | 32 | 2 |
| Proportion oligolectic: | | 5.2 | | 6.3 |

**Table S7 Notes.** The parasitic *L. lionotum* Sandhouse, *L. michiganense* Mitchell, and *L. platyparium* Robertson[54] do not collect pollen and so were excluded from the faunal list for this study. *Lasioglossum leucozonium* and *Lasioglossum zonulum* are introduced to North America[54,67] and so were also excluded.

**Table S8.** Summary of the species included in this study and the number of analyzed pollen loads per species.

| Michigan dataset | | British dataset | |
|---|---|---|---|
| Species | Sample size | Species | Sample size |
| *Andrena algida* | 11 | *Andrena alfkenella* | 23 |
| *Andrena aliciae* | 5 | *Andrena angustior* | 18 |
| *Andrena alleghaniensis* | 13 | *Andrena argentata* | 11 |
| *Andrena andrenoides* | 15 | *Andrena barbilabris* | 7 |
| *Andrena arabis* | 10 | *Andrena bicolor* | 41 |
| *Andrena barbilabris* | 23 | *Andrena bimaculata* | 13 |
| *Andrena bradleyi* | 9 | *Andrena chrysosceles* | 39 |
| *Andrena brevipalpis* | 25 | *Andrena cineraria* | 16 |
| *Andrena canadensis* | 6 | *Andrena denticulata* | 16 |
| *Andrena carlini* | 21 | *Andrena dorsata* | 55 |
| *Andrena carolina* | 5 | *Andrena flavipes* | 74 |
| *Andrena ceanothi* | 16 | *Andrena fucata* | 15 |
| *Andrena chromotricha* | 6 | *Andrena fulva* | 6 |
| *Andrena commoda* | 12 | *Andrena haemorrhoa* | 49 |
| *Andrena crataegi* | 28 | *Andrena helvola* | 12 |
| *Andrena cressonii* | 21 | *Andrena labialis* | 12 |
| *Andrena distans* | 5 | *Andrena labiata* | 10 |
| *Andrena erigeniae* | 7 | *Andrena minutula* | 74 |
| *Andrena erythrogaster* | 14 | *Andrena minutuloides* | 53 |
| *Andrena forbesii* | 25 | *Andrena nigriceps* | 5 |
| *Andrena frigida* | 11 | *Andrena nigroaenea* | 22 |
| *Andrena gardineri* | 5 | *Andrena nitida* | 28 |
| *Andrena geranii* | 5 | *Andrena scotica* | 39 |
| *Andrena helianthi* | 6 | *Andrena semilaevis* | 98 |
| *Andrena hippotes* | 22 | *Andrena subopaca* | 16 |
| *Andrena hirticincta* | 13 | *Andrena synadelpha* | 9 |
| *Andrena illinoiensis* | 8 | *Andrena thoracica* | 10 |
| *Andrena imitatrix* | 15 | *Andrena trimmerana* | 18 |
| *Andrena integra* | 10 | *Andrena wilkella* | 24 |
| *Andrena mandibularis* | 13 | | |
| *Andrena miranda* | 19 | plus 10 olioglectic species | |
| *Andrena miserabilis* | 23 | | |
| *Andrena nasonii* | 20 | *Andrena ferox* | |
| *Andrena nigrihirta* | 6 | *Andrena florea* | |
| *Andrena nivalis* | 13 | *Andrena fuscipes* | |
| *Andrena nubecula* | 7 | *Andrena hattorfiana* | |
| *Andrena nuda* | 10 | *Andrena marginata* | |
| *Andrena perplexa* | 10 | *Andrena nigrospina* | |
| *Andrena placata* | 9 | *Andrena nitidiuscula* | |
| *Andrena platyparia* | 7 | *Andrena niveata* | |
| *Andrena robertsonii* | 10 | *Andrena proxima* | |
| *Andrena rudbeckiae* | 8 | *Andrena tarsata* | |
| *Andrena rufosignata* | 19 | | |
| *Andrena rugosa* | 17 | | |
| *Andrena spiraeana* | 13 | | |
| *Andrena thaspii* | 9 | | |
| *Andrena vicina* | 37 | | |
| *Andrena virginiana* | 13 | | |
| *Andrena wellesleyana* | 9 | | |
| *Andrena w-scripta* | 16 | | |
| *Andrena ziziae* | 11 | | |

| | | | |
|---|---|---|---|
| Total | 671 | Total | 804 |
| *Lasioglossum acuminatum* | 20 | *Lasioglossum albipes* | 8 |
| *Lasioglossum anomalum* | 12 | *Lasioglossum calceatum* | 38 |
| *Lasioglossum cattellae* | 8 | *Lasioglossum fulvicorne* | 12 |
| *Lasioglossum cinctipes* | 12 | *Lasioglossum laevigatum* | 11 |
| *Lasioglossum coriaceum* | 24 | *Lasioglossum lativentre* | 18 |
| *Lasioglossum cressonii* | 28 | *Lasioglossum leucopus* | 11 |
| *Lasioglossum foxii* | 11 | *Lasioglossum leucozonium* | 26 |
| *Lasioglossum hitchensi* | 18 | *Lasioglossum malachurum* | 437 |
| *Lasioglossum imitatum* | 23 | *Lasioglossum morio* | 15 |
| *Lasioglossum laevissimum* | 19 | *Lasioglossum parvulum* | 12 |
| *Lasioglossum leucocomum* | 18 | *Lasioglossum pauxillum* | 70 |
| *Lasioglossum lineatulum* | 17 | *Lasioglossum prasinum* | 8 |
| *Lasioglossum lustrans* | 7 | *Lasioglossum puncticolle* | 13 |
| *Lasioglossum macoupinense* | 6 | *Lasioglossum villosulum* | 29 |
| *Lasioglossum nigroviride* | 7 | *Lasioglossum xanthopus* | 11 |
| *Lasioglossum oenothera* | 5 | *Lasioglossum zonulum* | 12 |
| *Lasioglossum paradmirandum* | 8 | | |
| *Lasioglossum paraforbesii* | 15 | | |
| *Lasioglossum pectorale* | 24 | | |
| *Lasioglossum pilosum* | 25 | | |
| *Lasioglossum quebecense* | 6 | | |
| *Lasioglossum subviridatum* | 8 | | |
| *Lasioglossum versans* | 7 | | |
| *Lasioglossum versatum* | 27 | | |
| Total | 355 | Total | 731 |
| Grand total | 1026 | Grand total | 1535 |

**Table S9: Pollen load data.** The pollen load percentages and botanical family counts of sampled *Andrena* and *Lasioglossum* bees from both the U.K. and Michigan, USA. Please find the data attached as an .xls file separated by location and botanical taxonomic level.

**Table S10: Simulation phenology parameters.** The phenology parameters for both pollinators and plants used in simulations. All potential combinations of the listed parameters were used to give 2072 unique phenological bases per network framework. Please find the data attached as an .xls file.
